# Brassinosteroid receptor BRL3 triggers systemic plant adaptation to elevated temperature from the phloem cells

**DOI:** 10.1101/2023.03.07.531487

**Authors:** Aditi Gupta, Andrés Rico-Medina, Fidel Lozano-Elena, Mar Marqués-Bueno, Juan B. Fontanet, Norma Fàbregas, Saleh Alseekh, Alisdair R. Fernie, Ana I. Caño-Delgado

## Abstract

Understanding plant receptor signaling is crucial to mitigate climate change impact on agriculture. BRs bind to membrane receptor-kinase BR-INSENSITIVE 1 (BRI1) in most plant cells that is essential to promote growth and stress responses, while the roles of vascular BRI1-LIKE1 and 3 (BRL3) receptors were considered redundant. While going unnoticed for twenty years, our study unveils that *brl3* mutants show conditional phenotypes to climate stress factors such as elevated temperatures, water deprivation and rising CO_2_ levels. In response to adverse climate conditions such as elevated temperature, BRL3 signaling at the phloem-companion cells can promote growth by activating BRI1-EMS-SUPPRESSOR1 (BES1) effector, hormonal homeostasis, and central carbon metabolism. This study shifts the paradigm for our present understanding of BR signaling and opens innovative strategies to develop climate-smart crops.

**One sentence summary:** Phloem-specific BRL3 receptor pathway controls plant adaption to elevated temperature.

## Main Text

The impact of climate change is posing a threat to agricultural and social sustainability on the planet^1, 2^. Indeed, stress factors such as extreme temperatures, drought, floods and rising atmospheric CO_2_ impact on sustainable food production on earth. Elevated temperatures provoked by CO_2_ emissions to the atmosphere have devastating effects on all agricultural ecosystems, bringing episodes of climate instability, water scarcity and more frequent outbreaks of pests and diseases, combining to result in dramatic crops loses every year^3–5^. Now, the search of innovative solutions to make climate-resilient crop toward the production of sustainable foods has become an emergency. Plants possess immense innate adaptive plasticity, and a more in-depth understanding of the underlying molecular mechanisms is crucial to strategize for sustaining populations under worsening climate change.

Plant receptor-like kinases (RLKs) initiate signaling cascades that control plant adaption to climate stress^6–8^. Brassinosteroid (BR) hormones directly bind to BR-INSENSITIVE 1 (BRI1) leucine rich repeat (LRR)-RLK family members on the plasma membrane^9–13^. Ligand perception triggers BRI1 to interact with the co-receptor BRI1 ASSOCIATED RECEPTOR KINASE 1 (BAK1)^14–16^ initiating a signaling cascade of phosphorylation events that control the expression of multiple BR-regulated genes mainly via the BRI1-EMS-SUPPRESSOR1 (BES1) and BRASSINAZOLE RESISTANT1 (BZR1) transcription factors^17–19^. Additionally, two BRI1-like receptors namely, BRI1-LIKE1 (BRL1) and BRL3 are mainly expressed in the plant vasculature^20^. While *bri1* mutant plants are dwarf and affected in most developmental responses along their life cycle, the absence of apparent phenotypes in *brl* mutants have kept the functional relevance of BRL receptors largely overlooked^20–24^. Our recent study reported that the overexpression of BRL3 receptors confers drought resistance without penalizing growth by promoting the accumulation of osmoprotectant compounds in response to drought^25^, suggesting stress specific roles for BRL3, yet awaited to be discovered.

Here, we postulated the hypothesis that plants use an alternate BR-mediated pathway through BRL3 receptors to promote growth adaptation in response to environmental signals^1, 26, 27^ . The *brl3* mutants (methods, 4 alleles, loss of function) exhibit wild-type like growth in favorable temperature (22°C) growth conditions (in agreement with^20^), yet a reduced hypocotyl length, due to reduced cell size, was observed when plants were exposed to elevated temperature (28°C) (Fig. 1a-d; Extended data Fig. S1). The conditional hypocotyl elongation defect of *brl3* mutants (Fig. 1c and d) was, however, restored by expressing native *pBRL3:BRL3-GFP* construct in the *brl3-2* mutant (*brl3* hereafter) background (Extended data Fig. S2). Further phenotypes associated to thermomorphogenesis such as petiole elongation and flowering time^28–30^ were also found to be perturbed in *brl3* mature plants (Fig. 1e and f; Extended data Fig. S3). By contrast, reduction of BRI1 signaling (*bri1* mutants or WT plants treated with BR-synthesis inhibitor BRZ220) led to reduced hypocotyl size irrespective of the temperature (Extended data Fig. S4 a-c). To rule-out the BRI1-dependency of the *brl3* phenotypes, we tested the transcript levels of BRI1, and other BRLs in the *brl3* mutant background, and found them to be at the WT-level (Extended data Fig. S5a). Together, our results unveil the existence of BR-mediated signaling cascade through BRL3 receptor to modulate growth adaptability under elevated ambient temperature conditions.

**Figure 1.**
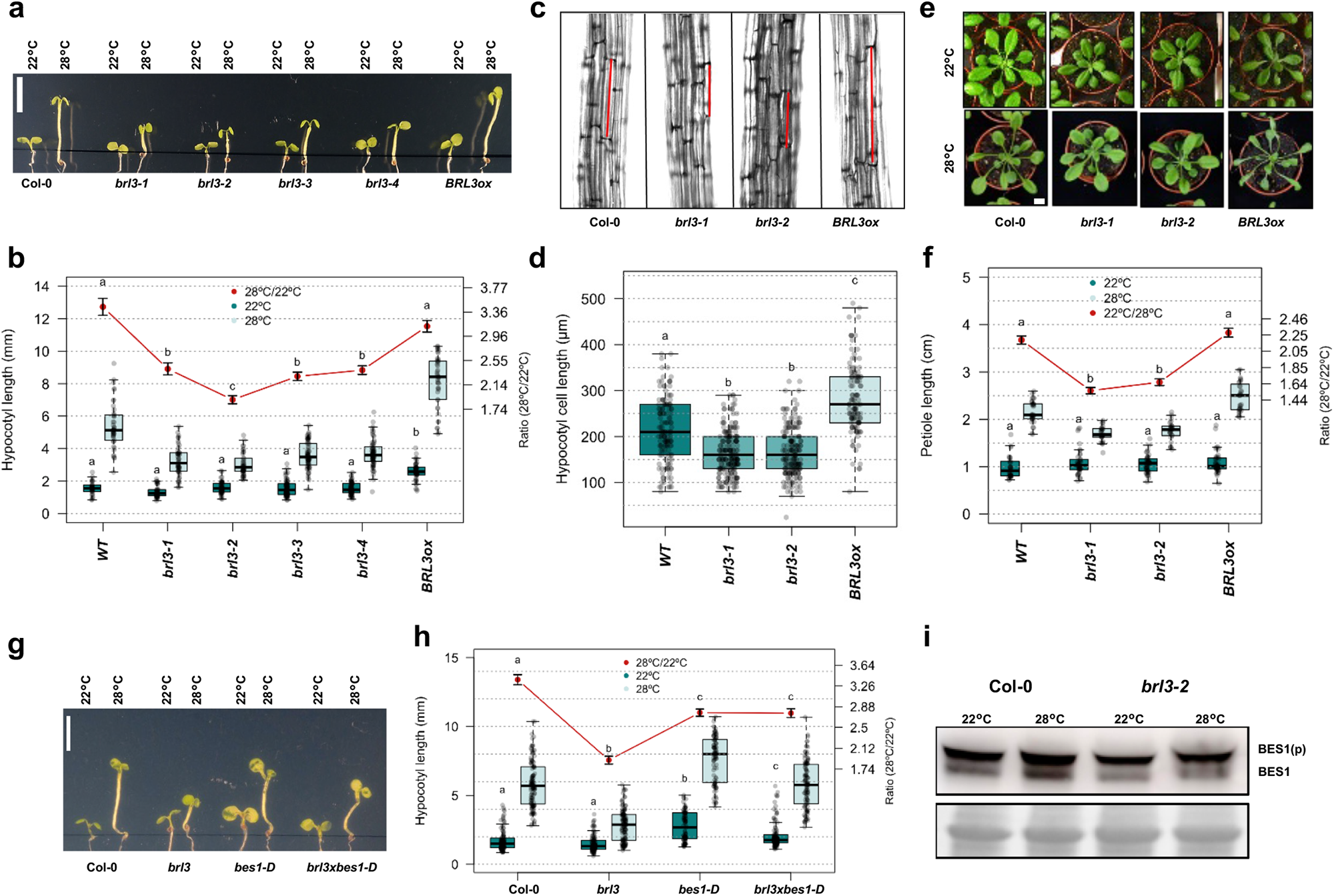
BRL3 receptor cascade controls thermomorphogenesis. **(a)** Pictures and **(b)** measurement of elevated temperature induced hypocotyl elongation growth of WT (Col-0), the *brl3-1*, *brl3-2*, *brl3-3* and *brl3-4* mutants, and *BRL3ox* seedlings grown at 22°C or 28°C for 6 days under LD conditions. Scale bar: 5 mm. **(c)** Pictures and **(d)** quantification of hypocotyl epidermal cell length from 6-day-old WT (Col-0), the brl3-1 and brl3-2 mutants and BRL3ox seedlings grown at 22°C or 28°C for 6 days under LD conditions. **(e)** Petiole elongation phenotypes and **(f)** quantification in 15-day-old WT (Col-0), the *brl3*-1 and *brl3*-2 mutants and *BRL3ox* plant grown at LD 22 °C condition and transferred to either 22°C or 28°C for 7-days. Scale bar:1 cm. **(g)** Pictures and **(h)** measurement of elevated temperature induced hypocotyl elongation growth of WT (Col-0), *brl3*-2, *bes1*-d and *brl3-2xbes1-d* seedlings. Scale bar: 5 mm. **(i)** Western blot showing BES1 phosphorylation status in WT (Col-0) and *brl3*-2 mutant seedlings at 22°C or 28°C. Five-days-old seedlings grown in LD at 22°C were transferred to wither LD at 22°C or LD at 28°C for 24h, BES1 was analyzed by Western blot hybridization with an anti-BES1 antibody. Boxplots depict the distribution of hypocotyl length (b, h), or Hypocotyl epidermal cell length (d) of 6-day-old seedlings, or petiole lengths (f) of rosette leaves from 22-day-old plants growing in control (dark green) or elevated temperature (light green) conditions. Red line depicts relative hypocotyl or petiole elongation upon high temperature (ratio 28 °C/22 °C ± s.e.m.). Different letters indicate a significant difference (p-value < 0.05) in a one-way ANOVA test plus Tukey’s HSD test. Boxplot represent the median and interquartile range (IQR). Whiskers depict Q1 – 1.5*IQR and Q3 + 1.5*IQR and points experimental observations. Data from four independent biological replicates (n > 50).

Upon elevated temperature, the core downstream components of BR-signaling cascades, BES1/BZR1 and BRASSINOSTEROID-INSENSITIVE 2 (BIN2), are known to interact with the PHYTOCHROME INTERACTING FACTOR 4 (PIF4)-mediated thermoresponsive pathway, driving the transcription of genes involved in growth adaptation^31–33^. The defective hypocotyl growth of *brl3* mutant was rescued by a gain-of-function *bes1-d* mutation (Fig. 1g and h). Strikingly, *brl3* mutants also displayed a reduced BES1 de-phosphorylation pattern upon elevated temperatures (Fig. 1i, Extended data Fig. S5b). Further, exogenous application of bikinin, an inhibitor of the BIN2 (BRI1-signaling negative regulator) rescued *brl3* conditional phenotypes on elevated temperature (Extended data Fig. S6) supporting the hypothesis that BRL3 uses the canonical BRI1 downstream signaling pathway to promote adaptive growth upon elevated temperatures.

Transcriptome analysis of *brl3* mutants uncovered that, at ambient temperature (22°C), BRL3-activated genes were enriched in growth, development, homeostasis, light response and tyrosine metabolism Gene Ontology (*GO*) categories, and BRL3-repressed genes were enriched in immune response, redox status and jasmonic acid-mediated response *GO* categories (|FC| > 1.5, FDR < 0.05; Supplementary Data S1, Extended data Fig. S7). The comparison of transcriptomic profiles of *brl3* at ambient and elevated (28°C) temperature (interaction term, see methods, Extended data Fig. S8a, Supplementary Data S2) revealed that genes involved in redox and abiotic stress responses (such as heat and water deprivation) failed to be activated the *brl3* mutant (under-response, Fig. 2a right). By contrast, negative regulators of ethylene and carboxylic acid and carbon catabolic processes responded stronger in the *brl3* mutant than in WT (over-response, Fig. 2a left). Among under-responsive genes annotated in abiotic stress responses (Fig. 2a), we found genes known to promote thermotolerance, such as *TEMPERATURE-INDUCED LIPOCALIN* (*TIL*), *HEAT SHOCK TRANSCRIPTION FACTOR A7A* (*HSFA7A*), and *MULTIPROTEIN BRIDGING FACTOR 1C* (*MBF1C*),^34–36^ and genes known to promote osmoprotectant synthesis such the *GALACTINOL SYNTHASE1* (*GolS1*)^37^ (Fig. 2b). These results agree with our previous report on the BRL3 overexpression lines (*BRL3ox)*^25^ . Further, core thermomorphogenesis genes such as those annotated as direct targets of BES1/BZR1, PIF4, AUXIN RESPONSE FACTOR 6 (ARF6) and HOMOLOG OF BEE2 INTERACTING WITH IBH 1 (HBI1) were also found to be significantly overrepresented in BRL3-regulated genes under elevated temperatures^38–42^ (Extended data Fig. S8b. These results suggest that BRL3 is necessary for the normal activation of high temperature response genes. Indeed, the *brl3* mutant seedlings were less adaptable to extreme temperatures (reduced survival) when subjected to a heat-shock treatment (150 min at 42°C) (Fig. 2c and d) and showed impaired response to osmotic stress in terms of hydrotropism (root growth reorientation towards water availability)^43^, and cell death in the root apex^44^ (Fig. 2 e-i). Conversely, *BRL3ox* were better adapted for both these traits^25^ (Fig. 2 e-i). When taken together, these data suggest that the BRL3 pathway is central for plant physiological and molecular adaption to both osmotic as well as high temperature stress.

**Figure 2.**
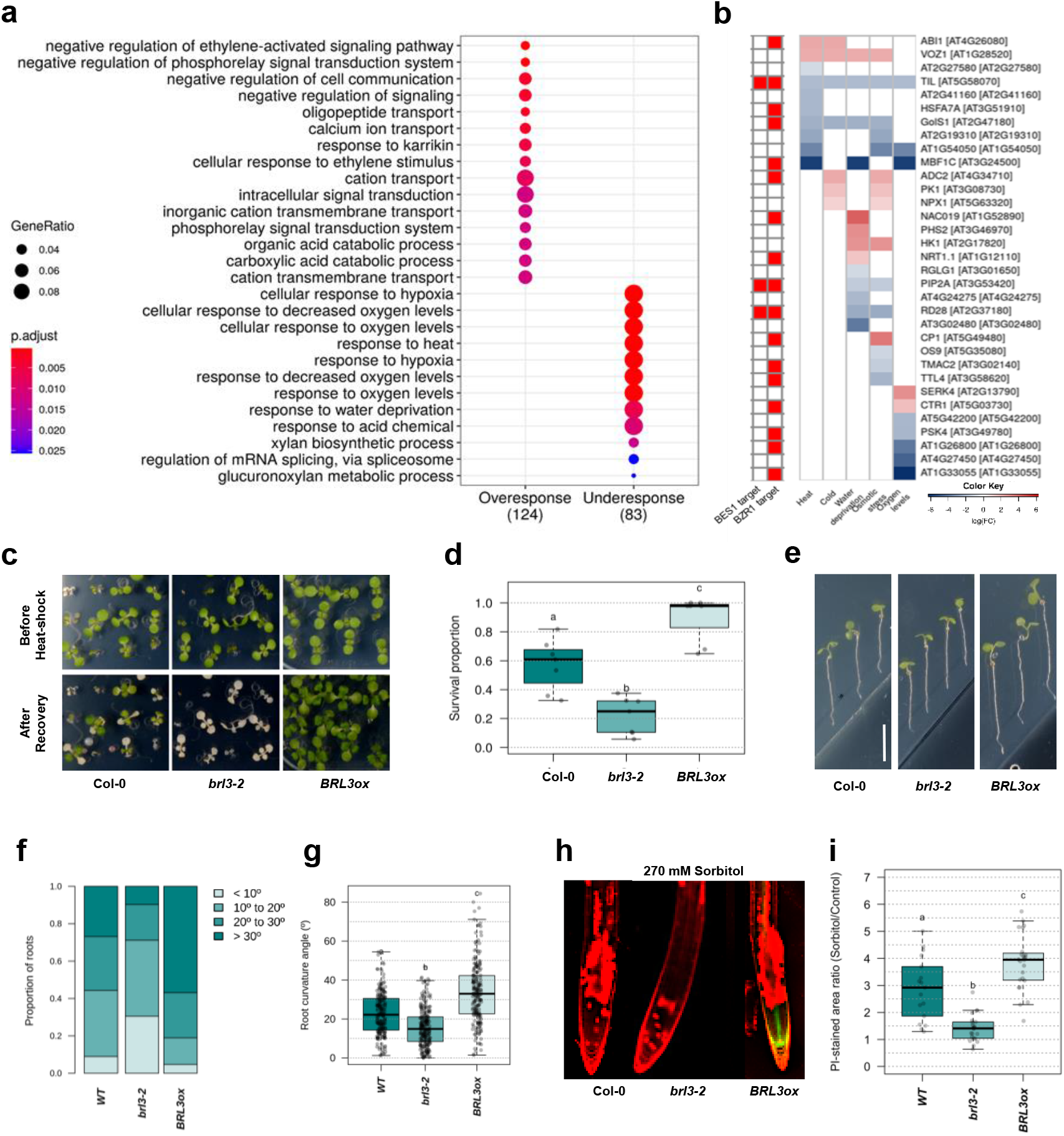
BRL3 is required for plant adaption to climate stresses. **(a)** GO Enrichment analysis based on genes affected by the interaction brl3∼temperature. Size of the node represents the number of annotated genes in a particular category and the color indicates its adjusted p-value upon enrichment test. **(b)** Deployment of genes affected by the interaction that are annotated as responding to abiotic stimulus [GO:0009628] and in its children categories (Responses to heat [GO:0009408], cold [GO:0009409], water deprivation [GO:0009414], osmotic stress [GO:0006970] and oxygen levels [GO:0070482]). Red means more high temperature-activation in brl3 mutant than in WT (putatively repressed by BRL3) and blue the opposite (putatively activated by BRL3). Small red boxes next to heatmap indicates that the gene is among the direct BES1 or BZR1 targets. **(c)** Pictures and **(d)** graphs showing plant survival after short-term heat-shock treatment (150 min at 42°C). Data from more than five independent biological replicates (n > 150). **(e)** Hydrotropic root curvature response in 6-day-old roots after 24 h of sorbitol-induced osmotic stress (270 mM). Scale bar: 10 mm. **(f)** Discrete distribution of root hydrotropic curvature angles in the different genotypes. **(g)** Continuous distribution of root curvature angles. Data from five independent biological replicates (n > 100). **(h)** Pictures showing cell death in root meristematic tissues after short term exposure to osmotic stress (8-10h), in WT, *brl3* and *BRL3ox* seedlings. Green channel (GFP) shows the localization of the BRL3 membrane protein receptor in the vascular tissues in primary roots. **(i)** Quantification of cell death in sorbitol-treated root tips. Boxplots show the relative PI staining (sorbitol/control) for each genotype. Averages from three independent biological replicates (n > 20). Different letters represent significant differences (p-value < 0.05) in an ANOVA plus Tukey’s HSD test. Boxplots represent the median and interquartile range (IQR). Whiskers depict Q1 – 1.5*IQR and Q3 + 1.5*IQR and points experimental observation.

Given that BRL3 receptors are natively expressed in phloem vascular tissues throughout the plant^20, 45^ (Extended data Fig. S9 a-g), we sought to investigate the cell-specific contribution of BRL3 signals to the observed phenotypes, by expressing BRL3 under different cell-type-specific promoters in the *brl3* mutant background (Fig. 3 a-g, Extended data fig. S9, S10). Driving BRL3 expression in the stele including xylem and phloem tissues (*pWOL:BRL3-GFP; brl3*) or specifically to the phloem companion cells (*pSUC2:BRL3-GFP; brl3*) rescued the thermomorphogenesis defects of *brl3* mutants (Fig. 3h and i, Extended data fig S9). Here, it is important to note that in the *pSUC2::BRL3-GFP; brl3* lines we enhanced the levels of BRL3 specifically in the phloem as compared to its native promoter driven expression in these cells (Extended data Fig. S9f-j). By contrast, BRL3 expression in either the epidermis (atrichoblast: *pGL2:BRL3-GFP; brl3* and trichoblast: *pEXP7:BRL3-GFP*; *brl3*), endodermis (*pSCR:BRL3-GFP; brl3*), root SCN (*pWOX5:BRL3-GFP; brl3*) or the meristem (*pRPS5A:BRL3-GFP; brl3*) failed to rescue the thermomorphogenic defects of the *brl3* mutants (Fig. 3 h and i; Extended data Fig S10c, d, e and g). Expressing BRL3 uniquely to other cells of the phloem tissue such as *pNAC86:BRL3-GFP; brl3* and *pCALS8:BRL3-GFP; brl3* was also able to rescue the thermomorphogenic defects of the *brl3* mutant (Extended data Fig S10a, b, e and f). This rescue was, however, partial due to weak expression strength of these promoters. It seems plausible that vascular cells of the phloem can modulate growth adaption via BRL3, independently of BRI1 (Extended data Fig S4a and b). Indeed, phloem specific expression of BRI1 could not rescue the hypocotyl elongation growth defects of the *brl3* mutant in our *pSUC2:BRI1-GFP; brl3* lines of Arabidopsis (Extended data Fig S11). Further, *pBRL3:BRL3-GUS* reporter analysis revealed that *BRL3* is spatially regulated upon elevated temperature *i.e.* induced in hypocotyl vasculature and repressed in the root vascular tissues (Fig. 3 j). Similar pattern was also observed for other vascular BR receptor *BRL1* expression (Fig. 3 j) suggesting that the BRLs receptors expression domain coincides with the organ and respective cellular region showing elongation growth responses upon elevated temperature. The epidermis has been proposed as a major site of action for generally controlling plant growth^46–48^. However, our data demonstrate that BRL3 receptor signaling modulates plant adaptive growth to elevated ambient temperature from the inner vascular cells of the phloem.

**Figure 3.**
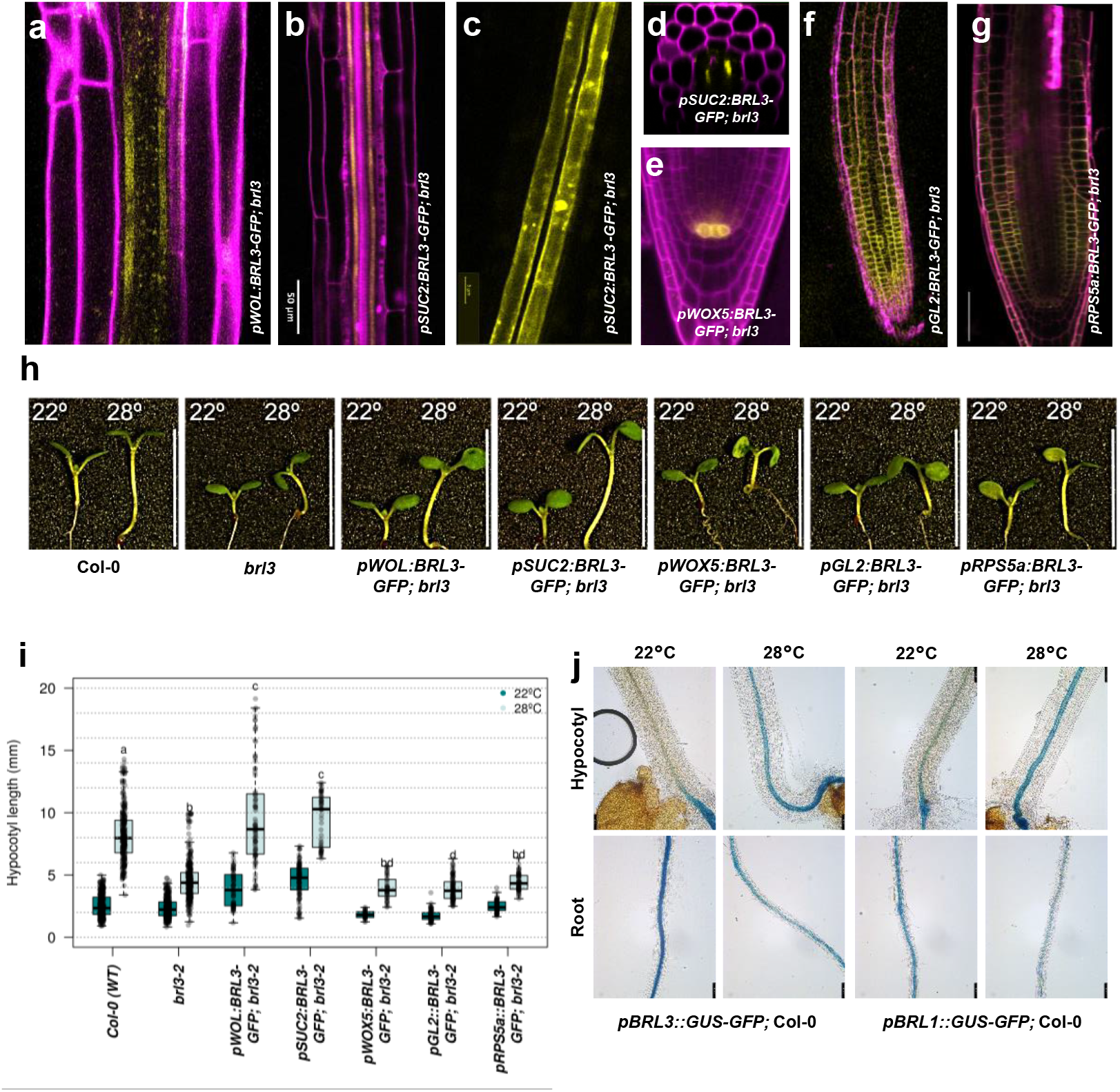
Local BRL3 receptor functions at the phloem companion cells. Confocal images of Arabidopsis roots showing BRL3 expression pattern in different cell/tissue types **(a)** Vasculature (*pWOL:BRL3-GFP;brl3)*; **(b-d)** phloem companion cells (*pSUC2:BRL3-GFP;brl3);* **(e)** root SCN (*pWOX5:BRL3-GFP;brl3)*; **(f)** epidermis (*pGL2:BRL3-GFP;brl3);* and **(g)** meristematic zone (*pRPS5a:BRL3-GFP;brl3).* **(h)** Pictures and **(i)** measurement of high temperature induced hypocotyl elongation growth of WT (Col-0), *brl3* mutant, and different cell/tissue specific complementation transgenic seedlings grown at 22 °C or 28 °C for 6 days under LD conditions. Scale bar: 10 mm. Boxplots depict the distribution of hypocotyl lengths in control (dark green) or elevated temperature (light green) conditions. Data from three independent biological replicates (n > 50). Boxplot represent the median and interquartile range (IQR). Whiskers depict Q1 – 1.5*IQR and Q3 + 1.5*IQR and points experimental observations. Different letters indicate a significant difference (p-value < 0.05) in a one-way ANOVA test plus Tukey’s HSD test. **(j)** Elevated temperature response of the BRL3 and BRL1 promoter activity. *pBRL3:GUS-GFP* and *pBRL1:GUS-GFP* seedlings were grown at 22 °C or 28 °C for 6 days under LD conditions followed by GUS staining. BRL3 promoter activity was induced in hypocotyls but reduced in roots upon elevated temperature.

Phloem function and its ability to transport resources is tightly controlled by the balance of carbon and water fluxes within the plant^49^. To elucidate phloem BRL3-driven transcriptional changes that contribute to temperature adaptation, we compared the transcriptomes of the *pSUC2:BRL3-GFP; brl3* line and *brl3* mutant shoots at ambient and elevated temperatures (Extended data Fig. S12, Supplementary Data S3, Fig. 4a, Supplementary Data S4). Genes with stronger response to temperature due to phloem specific-BRL3 expression were found to be enriched in energy generation processes (over-response, Fig. 4a left). These genes included genes associated with photorespiration as well as NADH-ubiquinone oxidoreductases and ATP syntheses linked to mitochondrial electron transfer chain, which in turn have great impact on redox homeostasis^50^ (*52*). By contrast, genes involved in auxin transport, (negative regulators of) ethylene signaling and steroid signaling, the later probably due to feedback mechanisms, displayed less deregulation in *pSUC2:BRL3; brl3* lines than in the *brl3* mutant subjected to increased temperatures (under-response, Fig. 4a right), suggesting partial rescue by BRL3 in the phloem. The mild repression of the photorespiration machinery observed in *pSUC2:BRL3-GFP; brl3* might result in better carbon utilization at elevated temperatures, as CO_2_/O_2_ selectivity of RuBisCO decreases under such conditions thereby inducing photorespiration pathway^51, 52^. Collectively, these results show significance of BRL3 signals from inner phloem companion cells in balancing plant adaptive growth by modulating plant energy status and hormone responses. We intersected deregulated genes in *brl3* (vs. WT) and *pSUC2:BRL3-GFP; brl3* (vs. *brl3*), and we found a moderate overlap in the response at 22°C (16%, Extended data fig S13) but a very high overlap in transcriptional responses at elevated temperatures (up to 60%), being 582 genes significantly deregulated in both comparisons and in opposite direction (“core genes”, Extended data fig S13, Supplementary Data S5). These comparisons support the increased relevance (and consistency) of BRL3 actions under elevated temperatures. We also confirmed an enrichment of BR-regulated genes^53, 54^ among the BRL3-regulated genes in both comparisons (*brl3* and *pSUC2:BRL3-GFP; brl3*) and “core genes” sets (Supplementary Data S6). Finally, in order to decipher the spatial distribution of (phloem) BRL3-induced transcriptional changes, we compared the deregulated transcripts with Arabidopsis leaves single cell expression atlas^55, 56^. We found that our deregulated genes were overrepresented with marker genes of vasculature and epidermis^55^, and specifically marker genes of vasculature subclusters as bundle sheath and xylem and phloem parenchyma^56^ (Supplementary Data S7). This reveals that most of the observed transcriptional response remains tissue-specific (or in close proximity) but part is translocated to distant tissues, although the mechanism remains unknown.

Since BRL3 from phloem companion cells regulates adaptive growth, we investigated whether BRL3 can affect phloem function. We used *pSUC2:GFP* reporter lines to check symplastic unloading of the free GFP signals in the roots^57, 58^. Plants obtained by the micrografting of *pSUC2:GFP* shoot scions onto *brl3* rootstocks displayed a significantly higher GFP signal at the root tip as compared to plants comprising of *pSUC2:GFP* shoot scions and a WT rootstock (Extended data fig S14a-c), indicating enhanced phloem unloading in the roots of *brl3* mutant. Further, in plants obtained by the micrografting of *pSUC2:GFP* shoot scions onto *pSUC2:BRL3-GFP;brl3* rootstocks displayed WT level phloem unloading in the roots (Extended data fig S14a-c). These observations suggest a role for BRL3 in nutrient and energy partitioning required for adaptive growth. To test this hypothesis, we performed metabolic profiling of WT, *brl3* and *pSUC2:BRL3-GFP*; *brl3* seedlings grown under ambient or elevated temperatures (Supplementary Data S8). The metabolic responses to elevated temperature were very conserved between WT and the *brl3* mutant shoots and roots, (Extended data Fig. S15). However, when comparing the metabolic footprint of shoot tissues of *brl3* (vs. WT) and *pSUC2:BRL3-GFP; brl3* (vs. *brl3*) at ambient temperature, tri-carboxylic acid (TCA) cycle intermediates, as well as the intimately associated metabolites glutamine, glutamic acid, alanine and pyruvic acid showed significantly altered levels in both comparisons (Fig. 4b), indicating that they were controlled by BRL3 action within the phloem. Strikingly, most of the changes were in the same direction (decreased in *brl3*) with the exception of pyruvic acid, which changes in an opposite direction (accumulated in *brl3* and decreased in *pSUC2:BRL3-GFP; brl3* shoots) (Fig. 4b). Pyruvic acid is a key central metabolite involved in several pathways of primary metabolism including glucose/gluconeogenesis and as an entry point into the TCA cycle. The accumulation of pyruvic acid in the mutant might indicate a poor carbon utilization (e.g. faster degradation of starch or stunted photosynthesis) and a slower TCA cycle resulting in a lower accumulation of intermediates (succinate, fumarate), reducing power and H^+^ gradient. On the other hand, decreased levels of pyruvate in *pSUC2:BRL3-GFP; brl3* plants could indicate a faster incorporation into TCA and increased flux, which would also result in a lower accumulation of intermediates (Fig. 4 c-g) but an increased H^+^ gradient and reducing power for mitochondrial respiration^59^. The combined evidence suggests that BRL3 signaling from vascular phloem cells contribute to boost central carbon metabolism, impacting the plant energy status and offering extra energy supply under challenging conditions. In addition, amino acid proline which functions as an osmoprotectant accumulated in plants expressing BRL3 solely in the phloem (Fig. 4 h), in agreement with the role of BRL3 in the activation of abiotic stress response genes observed in our transcriptomic data.

**Figure 4.**
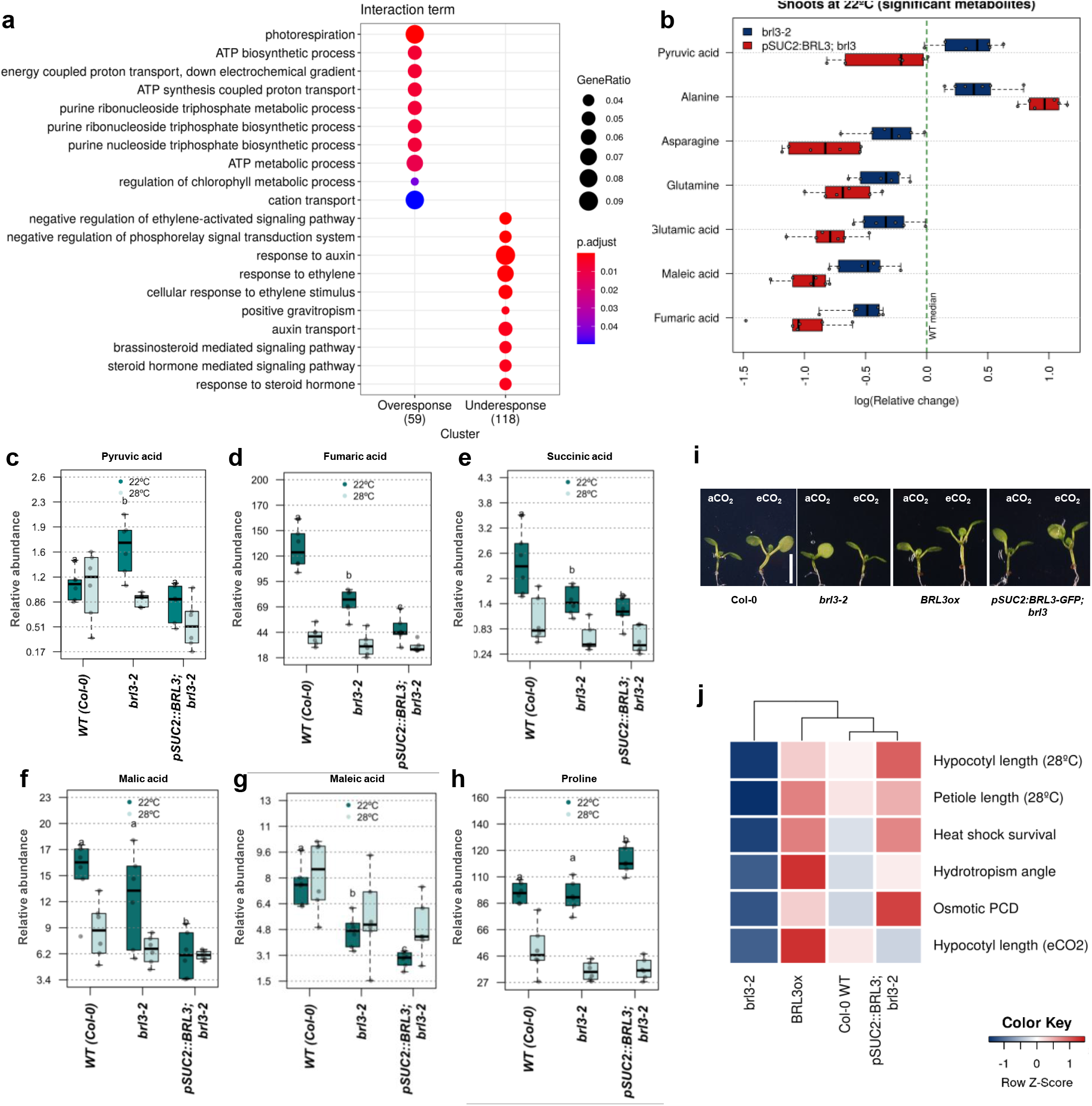
Phloem BRL3 signaling modulates carbon metabolism and energy flux to control adaptive growth. **(a)** GO Enrichment analysis based on genes affected by the interaction *pSUC2:BRL3-GFP; brl3*∼temperature i.e. genes that are either over-responding or under-responding to temperature in the *pSUC2:BRL3-GFP;brl3* line with respect the *brl3* mutant. Size of the node represents the number of annotated genes in a particular category and the color indicates its adjusted p-value upon enrichment test. **(b)** Metabolites differentially accumulated in shoots of 6-day-old seedlings of *brl3* vs. WT and *pSUC2:BRL3-GFP;brl3* vs. *brl3* at ambient temperature (22°C). Relative levels of **(c)** pyruvic acid and **(d-g)** intermediates of the Tricarboxylic Acid (TCA) cycle: Fumaric acid, Succinic acid, and configurational isomers Malic and Maleic acid, and **(h)** osmoprotectant amino acid proline in WT, *brl3* mutant and *pSUC2:BRL3-GFP;brl3* seedling shoot tissues at ambient (22°C) vs elevated (28°C) temperatures. Boxplots depict the distribution of indicated metabolite in control (dark green) or elevated temperature (light green) conditions. Different letters indicate a significant difference (p-value < 0.05) in a one-way ANOVA test plus Tukey’s HSD test. Data from six independent biological replicates. **(i)** Pictures showing hypocotyl elongation growth response to elevated CO_2_ (800 ppm) in 6-day-old seedlings of WT, *brl3, BRL3ox* and *pSUC2:BRL3-GFP;brl3* lines. Scale bar: 5 mm. **(j)** Stress traits matrix for all physiological assays performed on the roots and shoots of WT, *brl3, BRL3ox* and *pSUC2:BRL3-GFP;brl3* lines. Color bar depicts values for scaled data. In all the conditions tested, phloem specific BRL3 expression was able to rescue stress responsive defects of the *brl3* mutant.

Since BRL3 could potentially affect both photorespiration and carbon metabolism, we analyzed if BRL3 could modulate plant growth under elevated CO_2_ levels, a key climate factor contributing to global warming. In Arabidopsis, short-term exposure to high CO_2_ levels promotes the net rate of photosynthesis and hence stimulate overall plant growth^60, 61^. The *brl3* mutant was defective for hypocotyl elongation growth, whereas both the *pSUC2:BRL3-GFP; brl3* line and *BRL3ox* seedlings showed enhanced hypocotyl growth upon elevated CO_2_ (Fig 4 i, Extended data Fig. S16 a). Phloem-companion cell specific expression of BRL3 (*pSUC2:BRL3-GFP; brl3*) also rescued the defects in other stress adaptive responses such as heat and osmotic stress (Fig. 4 j, fig S16 b-g) indicating phloem-BRL3 signals are key for optimal plant growth and survival during climate change.

BRs are growth promoting plant hormones that play roles in abiotic stress resistance, including resistance against temperature extremes, high salt exposure and drought. While the stress-protective capacities of BRs have been known for decades, the molecular modes underlying BR function in abiotic stress responses had long remained elusive and are only starting to be revealed. After 20 years of its discovery and far from being redundant to BRI1^20–24^, our study uncovers a native role for BRL3 receptors in plant growth and adaption to adverse environmental conditions., i.e., such as elevated temperatures, elevated CO_2_, and water limitation.

At molecular level, similar to the BRI1, the BRL3 receptors bind brassinosteroid hormones with high affinity^20^ and can interact with known BRI1 interactors BAK1, BSK1 and BSK3^45^ suggesting common roles for the BRI1 and BRL3 receptors in the BR response pathway. The BRI1 receptors has also been shown to control plant response to abiotic stresses^62^, like the BRL3ox plants^25^. However, our study enlightens BRL3 roles on adaptive growth to the adverse environment, different of those of BRI1 receptor that constitutive in controlling growth regardless the climate conditions. This is also true for the majority Arabidopsis mutants in key regulatory pathways such as light, energy, sugar, auxin, GA, and ethylene etc, that are widely express throughout the plant, are deleterious even under optimal growth conditions^63–69^. Instead, BRL3 receptors discretely expressed in few cells, including those of the phloem^70^ pass inadvertence in normal conditions but is essential for adaptive growth responses to adverse environmental conditions specially elevated ambient temperature (Fig. 5). Noteworthily, micrografting data and the temperature regulation of BRL3 promoter expression in shoot and root vasculature and metabolite profiling suggest a key role of BRL3 in phloem function, resource partitioning as well as carbon metabolism. Phloem cells must generate some signal that relays to the epidermis and keeps the growth at a pace which allows optimal fitness during challenging conditions. In this regard, the identification of the intercellular signaling originating from the phloem cells towards the outer cell layers will be a priority in future research. Collectively, our data unveils that phloem specific BRL3 function is essential for the plant adaption to climate change and so must be the signals triggered by the BRL3 receptors within, thus shifting the paradigm of BR research and opening new avenues for engineering climate resilient crops.

**Figure 5.**
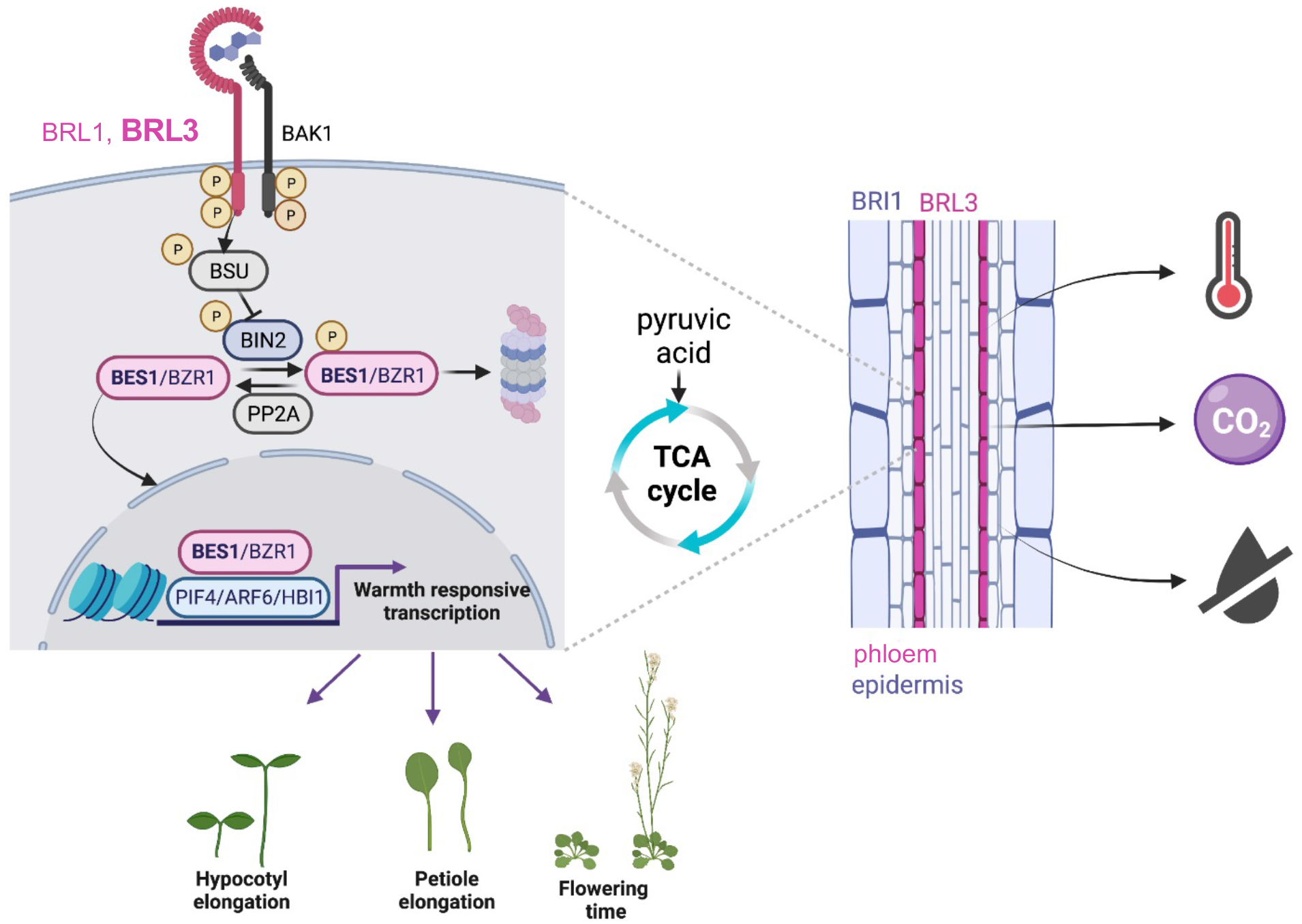
Vascular BR signaling via BRL3 regulates global transcriptome and plant carbon metabolism to control adaptive growth responses during adverse climate conditions such as increasing temperature, water limitation and elevated CO_2_ (Created with BioRender.com).

## Supporting information

Table S1. Primers used in this study

## Acknowledgments

We thank Y. Yin for the anti-AtBES1 antibody, and Centre Nacional d’Anàlisi Genòmica (CNAG-CRG) for support with RNA-seq.

## Funding

A.I.C.-D. has received funding from the European Research Council (ERC) under the European Union’s Horizon 2020 research and innovation programme (grant agreement 683163). A.I.C.-D. is a recipient of a BIO2016-78150-P grant funded by the Spanish Ministry of Economy and Competitiveness and Agencia Estatal de Investigación (MINECO/AEI) and Fondo Europeo de Desarrollo Regional (FEDER). A.G. has received funding by a postdoctoral fellowship from the “Severo Ochoa Programme for Centers of Excellence in R&D” 2016–2019 from the Ministerio de Ciencia e Innovación (SEV-2015-0533). A.G., F.L.-E., M.M.-B., A.R.M., and J.B.F. are funded by ERC2015-CoG–683163 granted to A.I.C.-D. F.L.-E and N.F. received funding from FEDER-BIO2016-78150-P and PIRSES-GA-2013-612583. A.R.-M. has received funding from Fundación Tatiana Pérez de Guzmán el Bueno. We acknowledge financial support from Spanish Ministry of Science and Innovation-State Research Agency (AEI) through the “Severo Ochoa Programme for Centres of Excellence in R&D” SEV-2015-0533 and CEX2019-000902-S and support from the CERCA Programme / Generalitat de Catalunya. S.A. and A.R.F. acknowledge funding of the PlantaSYST project by the European Union’s Horizon 2020 Research and Innovation Programme (SGA-CSA no. 664621 and no. 739582 under FPA no. 664620). The funders had no role in study design, data collection and analysis, decision to publish, or preparation of the manuscript.

## Author contributions

Conceptualization: ACD, AG

Methodology: AG, ARM, FLE, MM

Investigation: AG, FLE, MM, ARM, SA

Visualization: AG, FLE, MM, ARM, JBF

Funding acquisition: ACD, ARF

Project administration: ACD

Supervision: ACD

Writing – original draft: AG, ACD

Writing – review & editing: ACD, AG, FLE, MM, ARM, JBF, NF, SA, ARF

## Competing interests

The authors have filed a patent (EP 21 382 498.0.) on BRL3 function in climate adaption from phloem. They declare no other competing interests.

## Data and materials availability

All data are available in the main text or the supplementary materials.

## METHODS

### Plant materials and growth conditions

Wild-type Col-0 seedlings were used as a control (N1092: NASC (Nottingham Arabidopsis Stock Center number)). Four different T-DNA insertion mutant alleles of BRL3 were used *brl3-1*20 (*20*), *brl3-2* (SALK_006024C), *brl3-3* (SALK_079612) and *brl3-4* (SALK_111696) that are available in the ABRC (Arabidopsis Biological Resource Centre, Ohio State University) webpage. The *brl3-2* (SALK_006024C) loss-of-function allele was used for all the experiments described in this study unless specified. The seeds of *bri1-301, bak1-3, brl1, brl1brl3, brl1brl3bak1, bri1brl1brl3bak1, BRI1ox, BRL1ox, BRL3ox and bes1-d* were obtained as described25. Double mutants *bri1-301xbrl3* and *brl3xbes1-d* were created by making genetic cross between respective mutant alleles. Complementation line was created by making genetic cross between homozygous *pBRL3:BRL3-GFP* transgenic line^45^ and *brl3* mutant. Primers used for genotyping are listed in Extended Data Table S1.

*Arabidopsis thaliana* seeds were sterilized with 35% NaClO for 5 min and washed five times for 5 min with sterile dH2O. Sterilized seeds were stratified at 4 °C in the dark for 2–3 d. Seeds were placed on half strength Murashige and Skoog agar medium with vitamins and without sucrose (0.5MS-) solidified with 0.8% agar. Plates were transferred to chambers with controlled growth conditions (16 h of light/8 h of dark; 22 °C temperature, 60% relative humidity and 50– 70 μMol m−2 s−1 of cool-white fluorescent light).

Pharmacological treatments were performed in plates adding hormones/drugs in the media. Chemicals used include; 1µM BRZ220 (Gift from T. Nakano, RIKEN, Japan) or 50 µM Bikinin (200980, Sigma-Aldrich) for 6 days in control (long days, 22°C) or high temperature (long days, 28°C) conditions. Hypocotyl length were measured using the Image J software (v.1.48 v) (https://imagej.nih.gov/ij/).

### Phenotype analysis in seedlings

For root growth analysis, seedlings were grown in vertically placed 0.5MS-media plates (half strength MS media, with vitamins, without sugar solidified with 0.8% agar) in controlled growth conditions (Aralab 600; long days 16:8h day/night cycle; 22 °C). MyROOT software71 was used to compare root growth of plants. For thermomorphogenic hypocotyl elongation analysis, seedlings were grown in horizontally placed 0.5MS-media plates in controlled growth conditions (Aralab fitoclima600; long days 16:8h day/night cycle 60% relative humidity and 70 μmol m−2 s−1 of cool-white LEDs) at control (22 °C) or elevated (28 °C) temperature for 6 days. For Heat-shock assay, 7-day-old, homogenously grown seedlings were subjected to a 150 min heat shock at 42°C. Plants were moved back to control conditions for 5 days to check survival and recovery after heat-shock. For root hydrotropism, 5-day-old homogeneously grown seedlings were placed on 0.5MS-media plates in which lower part of the media was replaced with 0.5MS-media supplemented with 270 mM sorbitol to simulate a situation of reduced water availability and at 45° angle to avoid gravitropism effect. Root curvature angles were measured and analyzed 24h after using the Image J software. For cell death upon osmotic stress, 5-day-old seedlings grown on 0.5MS-media were shifted to either control or 270 mM sorbitol containing media for 8h, roots were stained with propidium iodide (10 μg/ml, PI, Sigma). PI stains the cell wall (control) and DNA in the nuclei upon cell death (sorbitol). Images were acquired with a confocal microscope (FV1000 Olympus). Cell death damage in primary roots was measured in a window of 500 µm from QC in the middle root longitudinal section (Image J). As an arbitrary setting to measure the stained area, a color threshold ranging from 160 to 255 in brightness was selected. To study the effect of elevated CO_2_ conditions on seedling growth, seedlings were grown in horizontally placed 0.5MS-media plates in controlled growth conditions (Aralab 600; long days 16:8h day/night cycle 60% relative humidity and 70 μmol m−2 s−1 of cool-white LEDs) at ambient CO_2_ (400 ppm) or elevated CO_2_ (800 ppm) conditions for 6-d. Hypocotyl length was measured using the ImageJ software (v.1.48 v) (https://imagej.nih.gov/ij/).

### Phenotype analysis in adult plants

Two-week-old seedlings grown in 0.5MS-agar plates were transferred individually to pots containing 30 ± 1 g of substrate (plus 1:8 v/v vermiculite and 1:8 v/v perlite) in normal growth conditions (long days, 22°C). For petiole elongation response to high temperatures, 2-week-old seedlings grown in MS1/2 agar plates were transferred individually to pots containing 30 ± 1 g of substrate (plus 1:8 v/v vermiculite and 1:8 v/v perlite). The plants were let to acclimatize for 2 days in normal growth conditions (long days, 22°C) before moving to either control (long days, 22°C) or high temperature (long days, 28°C) conditions. Petiole elongation growth in the longest rosette leaf was measured in 25d old plants using Image J software (v.1.48 v) (https://imagej.nih.gov/ij/). For flowering time analysis, plants were photographed and number of plants with >1 cm bolt were manually counted every day until all plants were bolted and had flowers.

### Plasmid construction and plant transformation

The pSUC2:BRL3-GFP, pWOL:BRL3-GFP, pWOX5:BRL3-GFP, pRPS5A:BRL3-GFP, pSCR:BRL3-GFP, pEXP7:BRL3-GFP, pGL2:BRL3-GFP, pNAC86:BRL3-GFP and pCALS8:BRL3-GFP and pSUC2:BRI1-GFP construct was generated using a recombination Gateway Multisite Cloning system (Invitrogen). Purified BRL3 and BRI1 gene PCR product was placed by directional cloning into the Gateway pDONR221® donor vector (12536-017; InvitrogenTM) by a recombination BP reaction. Gateway P4P1R vector containing cell specific promoters pSUC2, pWOL, pWOX5, pRPS5a, pGL2, pSCR, pEXP7 pCALS8 and pNAC86 were used as described in^72–74^. All the entry clones for cell/tissue specific promoters are available at http://www.ens-lyon.fr/DRP/SICE/Swelline.html. A recombination LR reaction was performed using the two sequenced pENTRY® vectors containing specific promoter and the BRL3 coding sequence were subcloned into the modified pDEST® gateway vector R4pGWB604 which contains a GFP tag and NPT gene for BASTA resistance. All inserts were fully sequenced to verify that no cloning errors occurred. The constructs were transformed into WT (Col-0) and *brl3* mutant (*brl3-*2) plants using *Agrobacterium tumefaciens* (GV3101)-mediated DNA transfer plants by floral dipping^75^. DNA rapid extraction protocol^76^ was used for all the plant genotyping experiments. Primers used for cloning, genotyping and sequencing are listed in Extended Data Table S1.

### GUS staining

*pBRL3:GFP-GUS* and *pBRL1:GFP-GUS* reporter lines^70^ were used to analyse the BRL3 and BRL1 expression pattern. The staining protocol was as follows: 6-day-old seedlings grown in ambient and elevated temperature conditions were submerged in ice-cold acetone 90% (v/v) and incubated for 20min on ice. Then acetone was rinsed twice with 1xPBS. Seedlings were incubated with GUS solution [100mM sodium phosphate buffer pH=7.2, 10mM sodium EDTA, 0.1% Triton X-100, 1m/ml 1.5-bromo-4-chloro-3-indolyl-beta-D-glucorince (Xglu; Duchefa, Harrlem, NL), 10mM potassium ferrocyanide and potassium ferricyanide] and incubated at 37°C overnight. Seedlings were rinsed with 1xPBS twice and let overnight in ethanol 70% (v/v). Seedlings were then mounted on slides and cover slides with chloral hydrate. Samples were observed at the following day under a stereomicroscope.

### RT-qPCR

RNA was extracted from the 15-day-old plants of different genotypes using the Plant Easy Mini Kit (Qiagen). The cDNA was obtained from RNA samples by using the Transcriptor First Strand cDNA Synthesis Kit (Roche) with oligo dT primers. The real-time qPCR reactions were performed from 10 ng of cDNA using LightCycler 480 SYBR Green I master mix (Roche) in 96-well plates according to the manufacturer’s recommendations. Ubiquitin (AT5G56150) was used as housekeeping gene for relativizing expression. Primers used are described in Extended Data Table S1.

### Western blot analysis

Five-day-old seedlings grown on 0.5MS-plates were collected directly or treated with 10 nM BL for 1 h, or kept in high temperature for 24 h. Forty entire seedlings of 5- or 6-day-old *Arabidopsis* seedlings were collected from each genotype and each treatment conditions, and ground into powder in liquid nitrogen. Protein extraction buffer (50 mM Tris-Cl, pH 6.8, SD, 5% glycerol) supplemented with 5% v/v beta mercaptoethanol and 1x protease inhibitor cocktail (Roche) was added to each sample followed by centrifugation at 18,000×g at 4 °C for 10 min. The resulting supernatants were transferred to a new microfuge tube. Eight µl of 6X Laemmli’s buffer was added to the protein samples to be loaded (40µl) and boiled at 95°C for 3 min. SDS/PAGE was performed to resolve the protein extracts. Loading of equal amounts of proteins was controlled by Ponceau-staining of the membranes. After electrophoresis, proteins were transferred to a PVDF (polyvinylidene difluoride) membrane (Millipore) using Wet transference method (Bio-Rad) and immune-detected with antibodies recognizing BES1^39^, Histone (NEB).

### Transcriptomic profiling analysis

For RNAseq analysis, shoot tissue was collected from 6-day-old seedlings grown under control (long days, 22°C) or high temperature (long days, 28°C) conditions. Two biological replicates were collected per genotype per experimental conditions. RNA was extracted with the Plant Easy Mini Kit (Qiagen) and quality checked using the Bioanalyser. Stranded cDNA libraries were prepared with TruSeq Stranded mRNA kit (Illumina). Paired-end sequencing, with 75-bp reads, was performed in an Illumina HiSeq500 sequencer, at a minimum depth of 32 M. Reads were trimmed 9 bp at their 3′ end and quality filtered using Trimmomatic. Remaining reads were mapped against the TAIR10 genome with “HISAT2”. Mapped reads were quantified at the gene level with “HtSeq” using Araport11 annotation file. Counts on chloroplast genes were eliminated prior analysis due to an identified artefact. Raw counts were TMM-normalized and differential expression calculated with “edgeR” package in R. To obtain genes with a different temperature response between genotypes, a lineal model accounting for the interaction temperature-genotype was applied. Values for the interaction term was evaluated in such comparisons. For pairwise comparisons, results were filtered for adjusted *p*-value (FDR) < 0.05 and FC > |1.5|. Raw reads and processed counts can be accessed at GEO with accession (GSE190107). Comparisons and enrichment testing with BR-regulated genes^53, 54^ were done by a two-sided Fisher’s exact tests. The same approach was taken with the gene markers of each tissue identified in the single cell experiments of^55, 56^.

### Metabolite profiling analyses

Seedlings were germinated in vertically placed 0.5MS-media plates for 6 days at control (22°C) or high (28°C) temperatures under long day growth conditions. Shoot and root tissue from six biological replicates were collected separately for both control and high temperature conditions, and for each genotype (WT, *brl3* and *pSUC2:BRL3-GFP;brl3*). Approximately 500 independent seedlings were bulked in each biological replicate. Frozen plant material was grinded in the Tissue Lyser Mixer-Mill (Qiagen) and were aliquoted into 50 mg samples for both shoot and root tissue samples (the exact weight was annotated for data normalization). Primary metabolite extraction was carried as follows (Lisec et al., 2006). One zirconia ball and 500 μl of 100% methanol premixed with ribitol (20:1) were added and samples were subsequently homogenized in the Tissue Lyser (Qiagen) 3 min at 25 Hz. Samples were centrifuged 10 min at 14,000 rpm (10 °C) and resulting supernatant was transferred into fresh tubes. Addition of 200 μl of CHCl_3_ and vortex ensuring one single phase followed by the addition of 600 μl of H2O and vortex 15 s. Samples were centrifuged 10 min at 14,000 rpm (10 °C). 100 μl from the upper phase (polar phase) were transferred into fresh eppendorf tubes (1.5 ml) and dried in the speed vacuum for at least 3 h without heating. 40 μl of derivatization agent (methoxyaminhydrochloride in pyridine) were added to each sample (20 mg/ml). Samples were shaken for 3 h at 900 rpm at 37 °C. Drops on the cover were shortly spun down. One sample vial with 1 ml MSTFA + 20 μl FAME mix was prepared. Addition of 70 μl MSTFA + FAMEs in each sample was done followed by shaking 30 min at 37 °C. Drops on the cover were shortly spun down. Samples were transferred into glass vials specific for injection in GC–TOF–MS. The GC–TOF–MS system comprised of a CTC CombiPAL autosampler, an Agilent 6890N gas chromatograph, and a LECO Pegasus III TOF–MS running in EI+ mode. Metabolites were identified by comparing to database entries of authentic standards (Kopka et al., 2005). Chromatograms were evaluated using Chroma TOF 1.0 (Leco) Pegasus software was used for peak identification and correction of RT. Mass spectra were evaluated using the TagFinder 4.0 software (Luedemann et al., 2008) for metabolite annotation and quantification (peak area measurements). The resulting data matrix was normalized using an internal standard, Ribitol, in 100% methanol (20:1), followed by normalization with the fresh weight of each sample. Overall metabolite levels were calculated summing up MS signals (non-normalized by fresh weight) in root and shoots, and normalizing by the sum of sample weights of shoot and root of the same replicate (overall sample weight). Pairwise comparisons between *pSUC2:BRL3-GFP; brl3* and *brl3* genotypes were performed through a two-tailed *t*-test, *p*-value < 0.05 (no multiple testing correction). For inferring metabolite transport between root and shoots, metabolite normalized measure in shoots was divided by overall metabolite level, yielding a matrix within the range [0-1], in which a value of 1 would mean all metabolite is accumulated in the shoot and 0 that all metabolite is accumulated in the roots. A value of 0.5 means equal distribution between shoot and roots. From metabolites whose overall levels are not changing (*t-test*, p-value > 0.05), pairwise comparisons between *pSUC2:BRL3-GFP; brl3* and *brl3* shoot/root ratio were performed to reveal differences in metabolite transport and/or accumulation (*t-test,* p-value < 0.05). All data transformation and calculations were performed in R.

### Micrografting and phloem unloading assay

Arabidopsis micrografting was performed on 7-d-old seedlings as described previously^77^ . Briefly, seeds were germinated and grown for 7-d under short day conditions. Homogenously grown seedlings were grafted and recovered on a water mounted filter paper/membrane at 24°C under long day conditions for 3-d. Successful grafts were transferred to fresh 0.5MS-media plates and grown for another 4-d at 22°C under long days conditions. GFP signals in the rootstocks were observed and images were acquired with a confocal microscope (FV1000 Olympus).

## EXTENDED DATA FIGURE LEGENDS

**Extended data figure S1. WT appearance of *brl3* mutant seedlings in normal growth conditions.**

**(a)** Diagram of *BRL3* showing the T-DNA insertion sites corresponding to different mutant alleles used in this study. **(b)** Relative *BRL3* transcript levels in different *brl3* mutant alleles. Transcript levels of *BRL3* were normalized to that of *UBIQUITIN* and are indicated as relative values, with that of the WT (Col-0) control set to 1. Data are presented as means ± SD calculated from three biological and technical replicates. Different letters represent significant differences (p-value < 0.05) in an ANOVA plus Tukey’s HSD test. **(c)** Pictures and **(d)** graph showing primary root growth of 7-d-old seedlings of WT (Col-0), and different *brl3* mutant alleles grown vertically in LD conditions at 22°C temperature. Scale bar: 5mm. Boxplots depict the distribution of primary root lengths. Boxplot represent the median and interquartile range (IQR). Whiskers depict Q1 – 1.5*IQR and Q3 + 1.5*IQR and points experimental observations. Data from three independent biological replicates (n > 50). Different letters represent significant differences (p-value < 0.05) in an ANOVA plus Tukey’s HSD test.

**Extended data figure S2. Complementation of thermomorphogenesis defect of *bparl3* mutant with native BRL3 expression.**

**(a)** Pictures and **(b)** measurement of high temperature induced hypocotyl elongation growth of WT (Col-0), *brl3-2* mutant and *pBRL3:BRL3-GFP; brl3* complementation lines grown at 22 °C or 28 °C for 6 days under LD conditions. Scale bar: 5mm. Boxplots depict the distribution of hypocotyl lengths in control (dark green) or elevated temperature (light green) conditions. Boxplot represent the median and interquartile range (IQR). Whiskers depict Q1 – 1.5*IQR and Q3 + 1.5*IQR and points experimental observations. Data from three independent biological replicates (n > 50). Different letters represent significant differences (p-value < 0.05) in an ANOVA plus Tukey’s HSD test.

**Extended data figure S3. Accelerated flowering-time phenotypes of *brl3* mutants.**

**(a)** Representative picture showing flowering phenotypes of the 25-d-old soil grown plants of WT, *brl3,* and *BRL3ox* lines. Scale bar: 5 cm. **(b)** Quantitative flowering time analysis (days to bolting) over a 35-days period in WT, *brl3,* and *BRL3ox* plants grown in control and elevated temperature conditions under long days in the greenhouse. Data from five independent biological replicates (*n* > 50).

**Extended data figure S4. Thermomorphogenesis phenotypes of mutants of BRI1-like receptors and co-receptor BAK1.**

**(a)** Pictures and **(b)** measurement of high temperature induced hypocotyl elongation growth of WT (Col-0), and single and higher order mutants as well as overexpressor lines of different components of BR receptor complex (BRI1, BRL1, BRL3, BAK1) grown at 22 °C or 28 °C for 6 days under LD conditions. Scale bar: 10mm. **(c)** measurement of hypocotyl elongation growth response of 6-d-old seedlings of WT (Col-0), *brl3-2* mutant and *BRL3ox* line grown at 22°C or 28°C for 6 days under LD conditions in media supplemented without or with Brassinazole (BRZ_220_, 1µM). Boxplots depict the distribution of hypocotyl lengths in control (dark green) or elevated temperature (light green) conditions. Boxplot represent the median and interquartile range (IQR). Whiskers depict Q1 – 1.5*IQR and Q3 + 1.5*IQR and points experimental observations. Data from three independent biological replicates (n > 50). Red line depicts relative hypocotyl elongation upon high temperature (ratio 28 °C/22 °C ± s.e.m.). Different letters represent significant differences (p-value < 0.05) in an ANOVA plus Tukey’s HSD test.

**Extended data figure S5. Involvement of other BR-signaling components in thermomorphogenesis phenotypes of *brl3* mutant.**

**(a)** Averaged and normalized transcript levels of BR receptors across conditions. The plot shows the log-transformed Reads Per Million (RPM, y-axis) of the BR receptors across RNAseq samples. **(b)** Measurement of phosphorylated vs unphosphorylated BES1 levels as obtained from WB analysis in WT (Col-0) and *brl3* mutant under normal vs elevated temperature conditions. Data are presented as means ± SD calculated from three biological and technical replicates. Asterisk represent significant differences (p-value < 0.05) in an Student’s paired t-test.

**Extended data figure S6. BIN2 acts downstream to BRL3 during thermomorphogenesis.**

**(a)** Pictures and **(b)** measurement of hypocotyl elongation growth response of 6-d-old seedlings of WT (Col-0) and *brl3-2* mutant grown at 22°C or 28°C for 6 days under LD conditions in media supplemented without or with Bikinin (50 µM). Scale bar: 5mm. Boxplots depict the distribution of hypocotyl lengths in control (dark green) or elevated temperature (light green) conditions in absence or presence of Bikinin. Boxplot represent the median and interquartile range (IQR). Whiskers depict Q1 – 1.5*IQR and Q3 + 1.5*IQR and points experimental observations. Data from three independent biological replicates (n > 50). Different letters represent significant differences (p-value < 0.05) in an ANOVA plus Tukey’s HSD test.

**Extended data figure S7. GO enrichment analysis of differentially regulated genes in *brl3-2* at 22°C in the RNAseq.**

Size of the node represents the number of annotated genes in a particular category and the color indicates its adjusted p-value upon enrichment test.

**Extended data figure S8. Transcriptome analysis of *brl3* mutants at high temperature.**

**(a)** Dotplot showing the comparison between transcript fold changes in WT (28°C vs. 22°C) and in brl3 (28°C vs. 22°C). Note that genes following the diagonal of the plot are responding equally. Red or blue dots represent genes that have been identified as affected by the interaction genotype∼temperature in the lineal model. **(b)** Enrichment in targets of core complex of cell elongation i.e. BAP module (BES1/BZR1, PIF4, ARF6) and HBI1 in genes affected by the interaction brl3-temperature.

**Extended data figure S9. BRL3 expression analysis and phenotype complementation from phloem companion cells.**

**(a-e)** GUS staining of 7-d-old *pBRL3::GFP-GUS;*Col-0 seedlings showing *BRL3* expression pattern in shoot apical meristem, cotyledon, hypocotyl and roots. Confocal imaging of roots maturation zone and root meristematic regions of 7-d-old **(f, g)** *pBRL3:BRL3-GFP;*Col-0 and **(h, i)** *pSUC2:BRL3-GFP;brl3* seedlings showing native and phloem-specific BRL3 expression respectively. **(j)** Quantification of mean phloem GFP intensity in phloem region in *pBRL3:BRL3-GFP;*Col-0 and *pSUC2:BRL3-GFP;brl3 roots* (Red box in f and g depicts the ROI used for measurements). Confocal images showing BRL3 expression pattern and vascular localization in **(k)** cotyledon and **(l)** root of *pSUC2:BRL3-GFP; brl3-2* transgenic seedlings. **(m)** Petiole elongation phenotypes and **(n)** quantification in WT (Col-0), *brl3-2* mutant and pSUC2:BRL3-GFP; brl3-2 transgenic plants grown at LD 22°C condition for 15-d and transferred to either 22°C or 28°C for 7-d. Boxplots depict the distribution of petiole lengths of rosette leaves from 22-d-old plants growing in control (dark green) or elevated temperature (light green) conditions. Different letters indicate a significant difference (p-value < 0.05) in a one-way ANOVA test plus Tukey’s HSD test. Boxplot represent the median and interquartile range (IQR). Whiskers depict Q1 – 1.5*IQR and Q3 + 1.5*IQR and points experimental observations. Data from three independent biological replicates (n > 30).

**Extended data figure S10. Local BRL3 expression in phloem can rescue *brl3* defects.**

Confocal imaging of 6-d-old **(a)** *pCALS8:BRL3-GFP;brl3* **(b)** *pNAC86:BRL3-GFP;brl3* **(c)** *pEXP7:BRL3-GFP;brl3* and **(d)** *pSCR:BRL3-GFP* seedling roots showing BRL3 expression in phloem pole pericycle, phloem sieve element, epidermis (trichoblasts) and endodermis respectively. **(e)** Pictures and (**f, g)** measurements of hypocotyl elongation response in WT (Col-0), *brl3* mutant, pCALS8:BRL3-GFP;brl3, pNAC86:BRL3-GFP;brl3, pEXP7:BRL3-GFP and pSCR:BRL3-GFP transgenic seedlings grown at 22 °C or 28 °C for 6 days under LD conditions. Scale bar: 10mm. Boxplots depict the distribution of hypocotyl lengths in control (dark green) or elevated temperature (light green) conditions. Boxplot represent the median and interquartile range (IQR). Whiskers depict Q1 – 1.5*IQR and Q3 + 1.5*IQR and points experimental observations. Red line depicts relative hypocotyl elongation upon high temperature (ratio 28 °C/22 °C ± s.e.m.). Data from three independent biological replicates (n > 50). Different letters represent significant differences (p-value < 0.05) in an ANOVA plus Tukey’s HSD test.

**Extended data figure S11. Local BRI1 expression from phloem companion cells can not rescue *brl3* thermomorphogenesis defects.**

**(a)**. Confocal imaging of primary roots maturation zone and meristematic regions of 7-d-old Arabidopsis *pSUC2:BRI1-GFP;brl3* seedlings. **(b)** Hypocotyl elongation phenotypes and **(c)** quantification in WT (Col-0), *brl3-2* mutant and *pSUC2:BRI1-GFP; brl3* transgenic grown at 22 °C or 28 °C for 6 days under LD conditions. Scale bar: 10mm. Boxplot represent the median and interquartile range (IQR). Whiskers depict Q1 – 1.5*IQR and Q3 + 1.5*IQR and points experimental observations. Red line depicts relative hypocotyl elongation upon high temperature (ratio 28 °C/22 °C ± s.e.m.). Data from three independent biological replicates (n > 30). Different letters represent significant differences (p-value < 0.05) in an ANOVA plus Tukey’s HSD test.

**Extended data figure S12. GO enrichment analysis of differentially regulated genes upon phloem specific BRL3 expression.**

GO enrichment analysis of genes up or downregulated in *pSUC2:BRL3-GFP; brl3-2* vs. *brl3-2* at ambient temperature (22°C) conditions. Size of the node represents the number of annotated genes in a particular category and the color indicates its adjusted p-value upon enrichment test.

**Extended data figure S13. “Core Genes”. Genes deregulated in both comparisons, *brl3* vs. WT and *pSUC2:BRL3; brl3* vs. *brl3,* and in opposite direction.**

**(a, b)** Overlapping of the transcriptional responses in the comparisons *brl3-2* vs. WT and pSUC2:BRL3; *brl3* vs. *brl3* at 22°C (a) and 28°C (b). Note the increased overlap at elevated temperature, up to 60% of DEG in *brl3*. **(c, d)** Scatter plots showing fold changes of both comparison (x-y axis) at 22°C (c) and 28°C (d). Genes following the inverse diagonal (purple) show an opposite behavior in *brl3* than in pSUC2:BRL3. These have high probabilities to be directly BRL3-regulated and we denominated them “Core genes”. **(e, f)** GO enrichment analysis of the core genes, separating the enriched categories among the BRL3-activated genes (red) and BRL3-repressed genes (blue), either at 22°C (e) and 28°C (f).

**Extended data figure S14. Micrografting experiments showing BRL3 effect on phloem function.**

**(a)** Graphical representation of the symplastic unloading of free-GFP using micrografting between *pSUC2:GFP* scion and rootstock from either WT Col-0 or *brl3* mutant seedlings. **(b)** Live confocal imaging of 6-day-post-grafting Arabidopsis root tips. Boxes account for the ROI area where GFP was measured, using the same ROI for all images placed right above the QC. In the labels, upper line indicates upper part of the chimeric plant after grafting (cotyledons and upper part of the hypocotyl), lower line indicates root part of the chimeric plant (upper part of the hypocotyl and primary root). **(c)** Mean GFP intensity in the selected area from B. Notice an increased unloading in root tip in brl3 root. Letters depict significant differences following ANOVA with Tukey Post-Hoc HSD test. data from three biological replicates, 9<n≤15.

**Extended data figure S15. Metabolomics analysis of Col-0, *brl3* and pSUC2BRL3 under elevated temperatures**

Schematic representation of the specific metabolic response of **(a)** shoots and **(b)** roots of 6-d-old seedlings of WT (Col-0), brl3 and *pSUC2:BRL3-GFp;brl3* lines exposed to elevated temperature (28 °C) versus optimal temperature (28 °C) under LD conditions. Colored cells represent the log fold-changes. Statistically significant changes (28°C vs 22°C) are denoted by thick black borders on the boxes. Asterisks denote metabolites which have a statistically significant 28°C/22°C ratio between genotypes. Note that most of the metabolites respond similarly to temperature among genotypes.

**Extended data figure S16. BRL3 signals from phloem regulate plant adaptation to multiple climate stress.**

**(a)** Graphs showing hypocotyl elongation growth response to elevated CO_2_ in 6-d-old seedlings of WT, *brl3* and *pSUC2:BRL3-GFP;brl3* lines. **(b)** Pictures showing plant survival phenotypes after short-term heat-shock treatment (150 min at 42°C), **(c)** Quantification of survival rates upon heat-shock treatments. **(d)** Root hydrotropism response, **(e)** Distribution of the root angles upon hydrotropism response. **(f)** Cell death in root meristematic tissues after short term exposure to osmotic stress (8-10h), in WT, *brl3* and *pSUC2:BRL3-GFP;brl3* seedlings. **(g)** Quantification of cell damage due osmotic stress. As PI-stained ratio between treated and untreated roots. In all the conditions tested, phloem specific BRL3 expression was able to rescue stress responsive defects of the *brl3* mutant.

**Extended Data Table S1.** List of primers

## SUPPLEMENTARY DATA TABLES

**Supplementary Data S1.** List of deregulated genes WT vs *brl3* at 22°C

**Supplementary Data S2.** List of genes with a different temperature response between *brl3* and WT (genotype*treatment interaction-affected genes)

**Supplementary Data S3.** List of deregulated genes *pSUC2:BRL3-GFP; brl3* vs *brl3* at 22°C

**Supplementary Data S4.** List of genes with a different temperature response between *pSUC2:BRL3-GFP;brl3* and *brl3* (genotype*treatment interaction-affected genes)

**Supplementary Data S5**. List of Differentially Expressed Genes (DEGs) overlapping between comparisons *brl3* vs. WT and p*SUC2:BRL3-GFP;brl3* vs *brl3* at 22°C and 28°C in opposite direction (“core genes”)

**Supplementary Data S6.** BR-regulated genes that are BRL3-regulated from the phloem

**Supplementary Data S7.** Values of over-representation tests for tissue-specific markers (as for single cell transcriptomics) among DEGs and “core genes”, and the vascular-specific deregulated marker genes.

**Supplementary Data S8.** List of metabolites

