## Extended Data Figures S1-S16 for "Brassinosteroid receptor BRL3 triggers systemic plant adaptation to elevated temperature from the phloem cells"

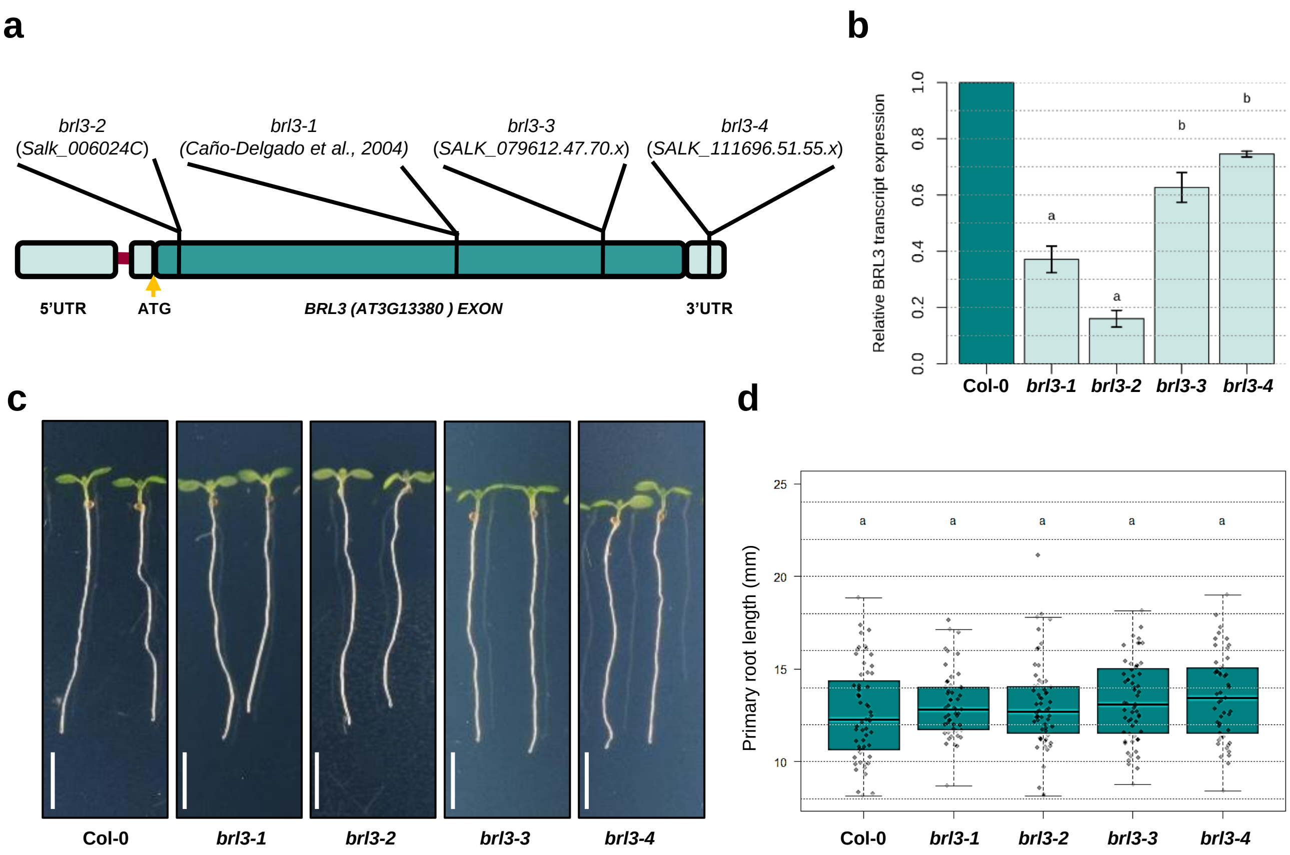


**Extended data figure S1. WT appearance of *brl3* mutant seedlings in normal growth conditions.** **(a)** Diagram of *BRL3* showing the T-DNA insertion sites corresponding to different mutant alleles used in this study. **(b)** Relative *BRL3* transcript levels in different *brl3* mutant alleles. Transcript levels of *BRL3* were normalized to that of *UBIQUITIN* and are indicated as relative values, with that of the WT (Col-0) control set to 1. Data are presented as means ± SD calculated from three biological and technical replicates. Different letters represent significant differences ($p$-value < 0.05) in an ANOVA plus Tukey's HSD test. **(c)** Pictures and **(d)** graph showing primary root growth of 7-d-old seedlings of WT (Col-0), and different *brl3* mutant alleles grown vertically in LD conditions at 22°C temperature. Scale bar: 5mm. Boxplots depict the distribution of primary root lengths. Boxplot represent the median and interquartile range (IQR). Whiskers depict Q1 – 1.5*IQR and Q3 + 1.5*IQR and points experimental observations. Data from three independent biological replicates (n > 50). Different letters represent significant differences ($p$-value < 0.05) in an ANOVA plus Tukey's HSD test.

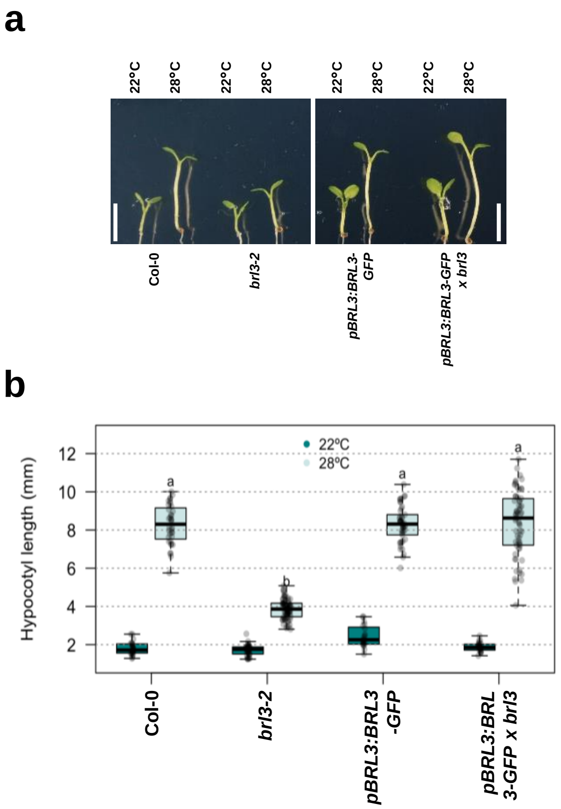


**Extended data figure S2. Complementation of thermomorphogenesis defect of *brl3* mutant with native BRL3 expression.**

**(a)** Pictures and **(b)** measurement of high temperature induced hypocotyl elongation growth of WT (Col-0), *brl3-2* mutant and *pBRL3:BRL3-GFP; brl3* complementation lines grown at 22 °C or 28 °C for 6 days under LD conditions. Scale bar: 5mm. Boxplots depict the distribution of hypocotyl lengths in control (dark green) or elevated temperature (light green) conditions. Boxplot represent the median and interquartile range (IQR). Whiskers depict Q1 – 1.5*IQR and Q3 + 1.5*IQR and points experimental observations. Data from three independent biological replicates ($n > 50$). Different letters represent significant differences ($p$-value < 0.05) in an ANOVA plus Tukey's HSD test.

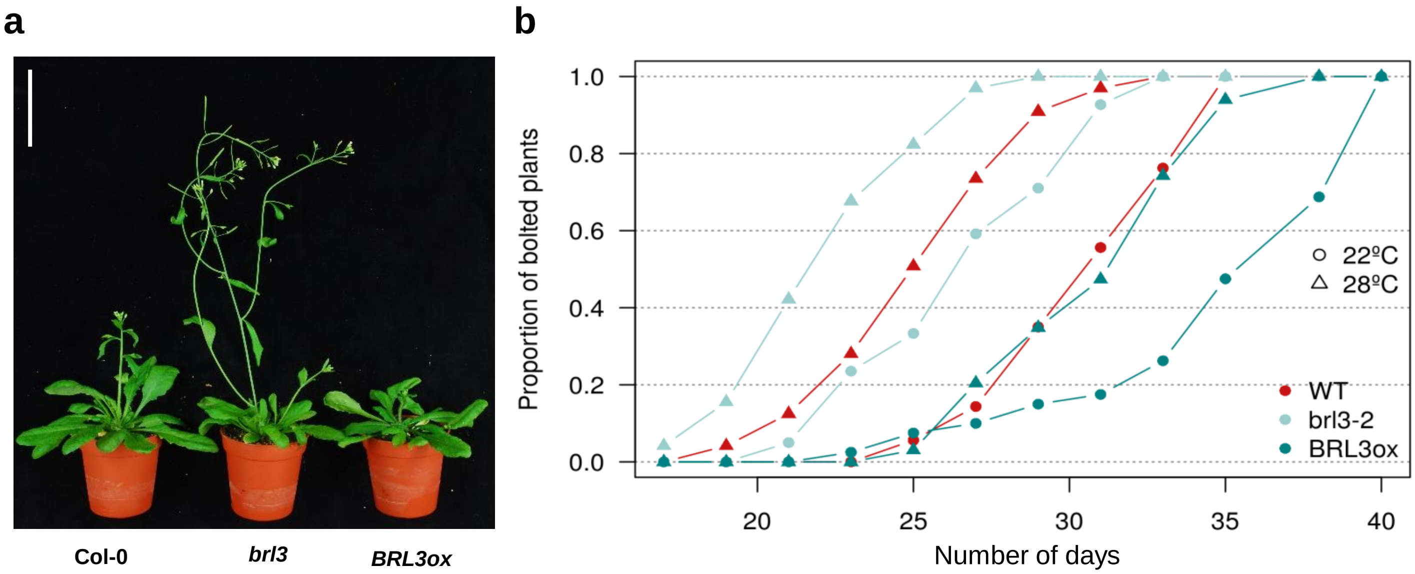


**Extended data figure S3. Accelerated flowering-time phenotypes of *brl3* mutants.**
**(a)** Representative picture showing flowering phenotypes of the 25-d-old soil grown plants of WT, *brl3,* and *BRL3ox* lines*.* Scale bar: 5 cm. **(b)** Quantitative flowering time analysis (days to bolting) over a 35-days period in WT, *brl3,* and *BRL3ox* plants grown in control and elevated temperature conditions under long days in the greenhouse. Data from five independent biological replicates ($n > 50$).

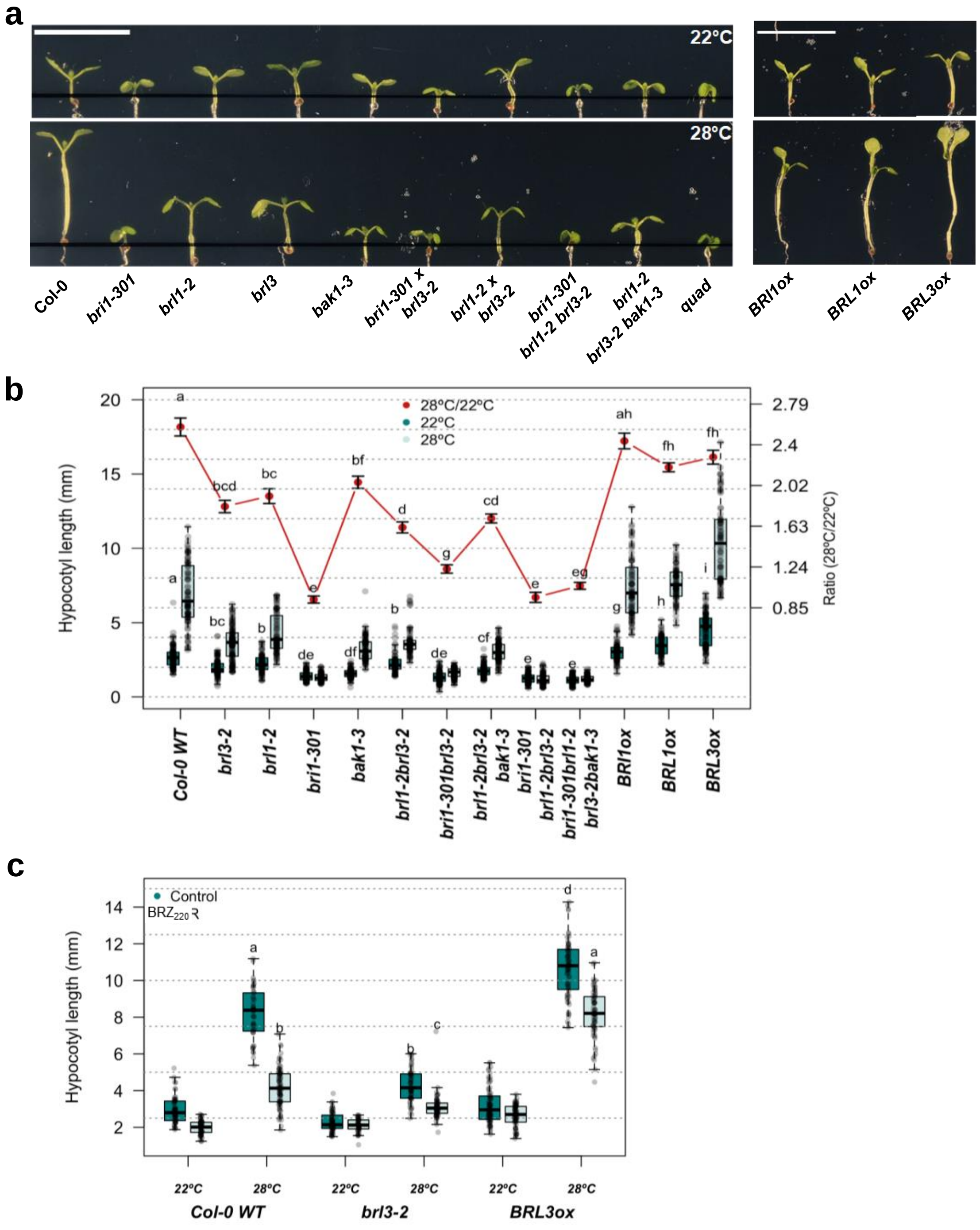


**Extended data figure S4. Thermomorphogenesis phenotypes of mutants of BRI1-like receptors and co-receptor BAK1.**

**(a)** Pictures and **(b)** measurement of high temperature induced hypocotyl elongation growth of WT (Col-0), and single and higher order mutants as well as overexpressor lines of different components of BR receptor complex (BRI1, BRL1, BRL3, BAK1) grown at 22 °C or 28 °C for 6 days under LD conditions. Scale bar: 10mm. **(c)** measurement of hypocotyl elongation growth response of 6-d-old seedlings of WT (Col-0), *brl3-2* mutant and *BRL3ox* line grown at 22°C or 28°C for 6 days under LD conditions in media supplemented without or with Brassinazole ($BRZ_{220}$, 1µM). Boxplots depict the distribution of hypocotyl lengths in control (dark green) or elevated temperature (light green) conditions. Boxplot represent the median and interquartile range (IQR). Whiskers depict Q1 – 1.5*IQR and Q3 + 1.5*IQR and points experimental observations. Data from three independent biological replicates ($n > 50$). Red line depicts relative hypocotyl elongation upon high temperature (ratio 28 °C/22 °C ± s.e.m.). Different letters represent significant differences ($p$-value < 0.05) in an ANOVA plus Tukey's HSD test.

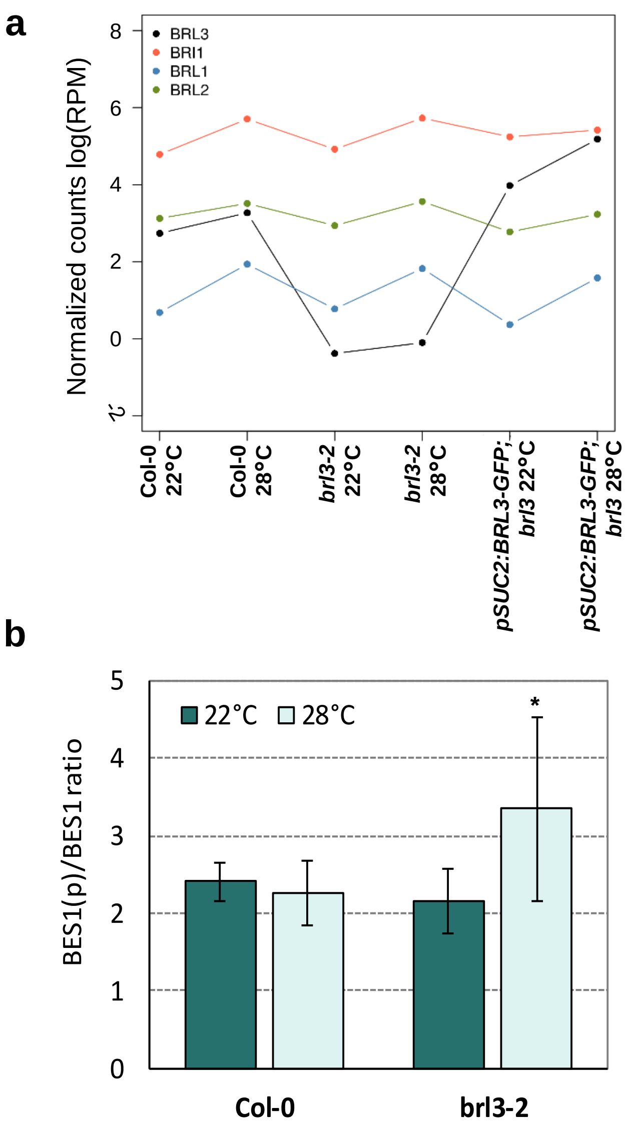


**Extended data figure S5. Involvement of other BR-signaling components in thermomorphogenesis phenotypes of brl3 mutant.**
**(a)** Averaged and normalized transcript levels of BR receptors across conditions. The plot shows the log-transformed Reads Per Million (RPM, y-axis) of the BR receptors across RNAseq samples. **(b)** Measurement of phosphorylated vs unphosphorylated BES1 levels as obtained from WB analysis in WT (Col-0) and *brl3* mutant under normal vs elevated temperature conditions. Data are presented as means ± SD calculated from three biological and technical replicates. Asterisk represent significant differences (p-value < 0.05) in an Student's paired t-test.

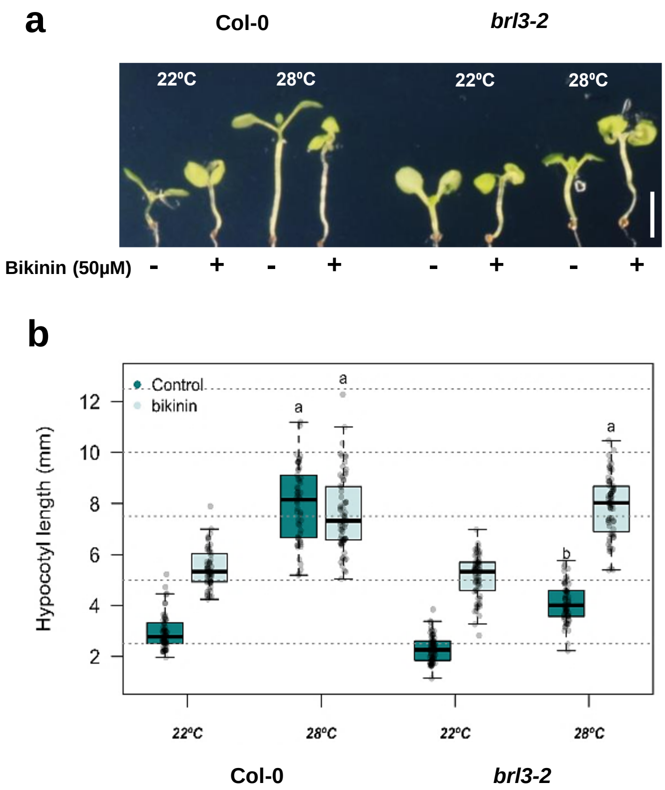


**Extended data figure S6. BIN2 acts downstream to BRL3 during thermomorphogenesis.**

**(a)** Pictures and **(b)** measurement of hypocotyl elongation growth response of 6-d-old seedlings of WT (Col-0) and *brl3-2* mutant grown at 22°C or 28°C for 6 days under LD conditions in media supplemented without or with Bikinin (50 µM). Scale bar: 5mm. Boxplots depict the distribution of hypocotyl lengths in control (dark green) or elevated temperature (light green) conditions in absence or presence of Bikinin. Boxplot represent the median and interquartile range (IQR). Whiskers depict Q1 – 1.5*IQR and Q3 + 1.5*IQR and points experimental observations. Data from three independent biological replicates ($n > 50$). Different letters represent significant differences ($p$-value $< 0.05$) in an ANOVA plus Tukey's HSD test.

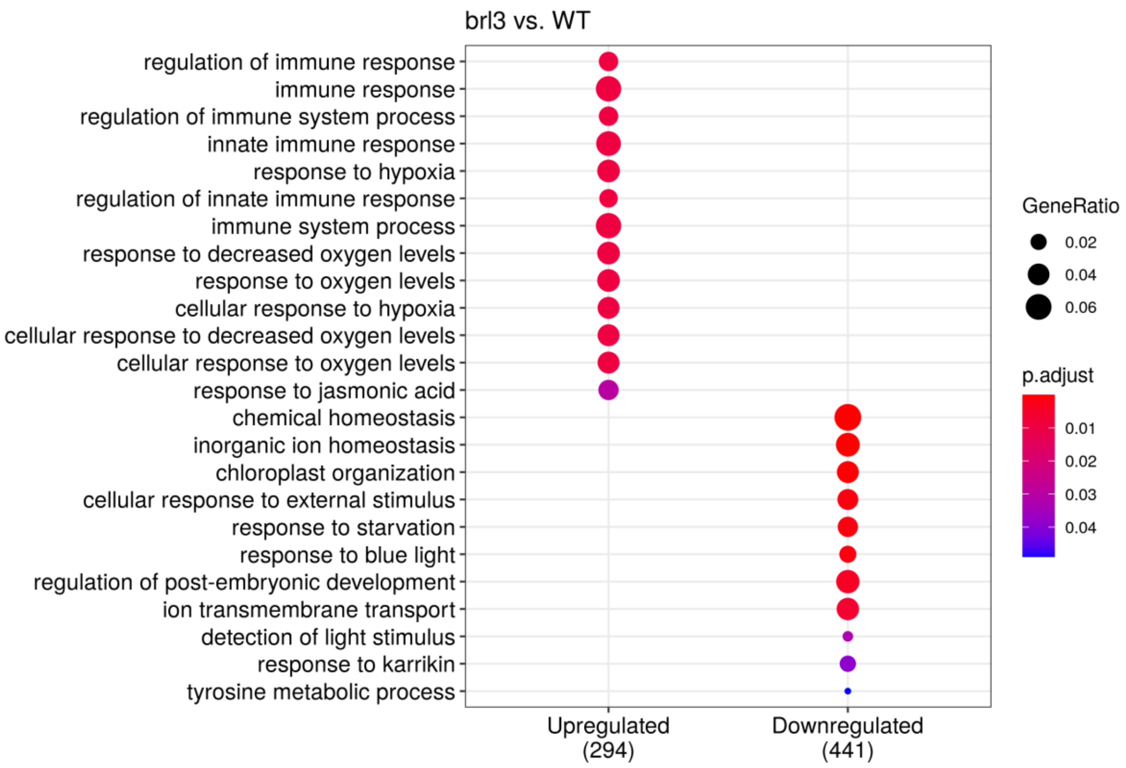


**Extended data figure S7. GO enrichment analysis of differentially regulated genes in *brl3-2* at 22ºC in the RNAseq.**

Size of the node represents the number of annotated genes in a particular category and the color indicates its adjusted p-value upon enrichment test.

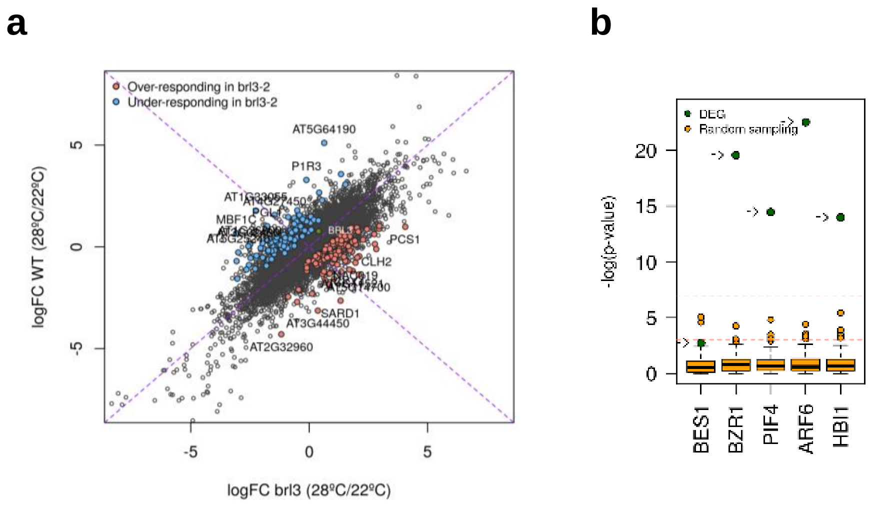


**Extended data figure S8. Transcriptome analysis of *brl3* mutants at high temperature.**

**(a)** Dotplot showing the comparison between transcript fold changes in WT (28ºC vs. 22ºC) and in brl3 (28ºC vs. 22ºC). Note that genes following the diagonal of the plot are responding equally. Red or blue dots represent genes that have been identified as affected by the interaction genotype~temperature in the lineal model. **(b)** Enrichment in targets of core complex of cell elongation i.e. BAP module (BES1/BZR1, PIF4, ARF6) and HBI1 in genes affected by the interaction brl3-temperature.

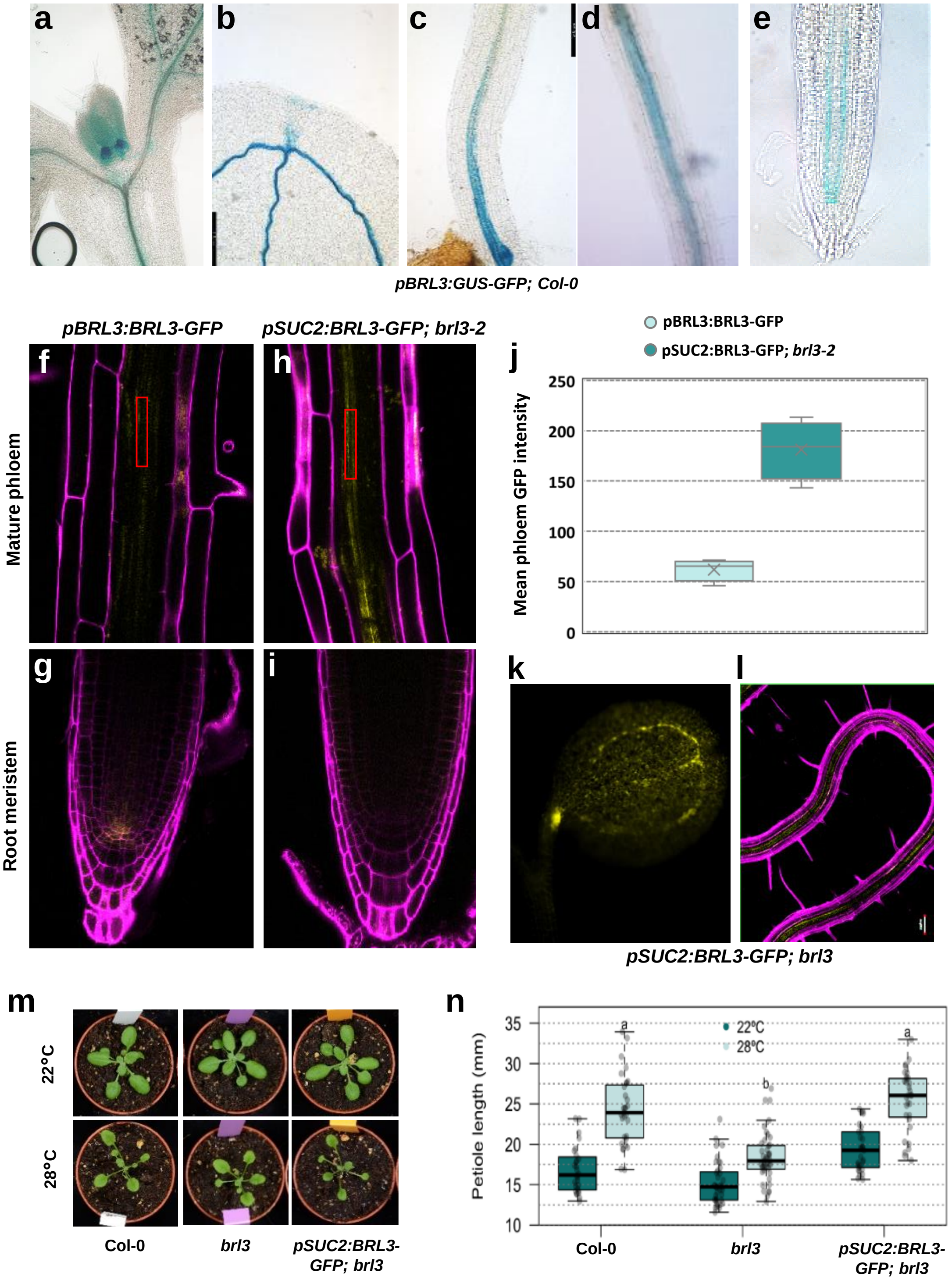


**Extended data figure S9. BRL3 expression analysis and phenotype complementation from phloem companion cells.** **(a-e)** GUS staining of 7-d-old *pBRL3::GFP-GUS;*Col-0 seedlings showing *BRL3* expression pattern in shoot apical meristem, cotyledon, hypocotyl and roots*.* Confocal imaging of roots maturation zone and root meristematic regions of 7-d-old **(f, g)** *pBRL3:BRL3-GFP;*Col-0 and **(h, i)** *pSUC2:BRL3-GFP;brl3* seedlings showing native and phloem-specific BRL3 expression respectively. **(j)** Quantification of mean phloem GFP intensity in phloem region in *pBRL3:BRL3-GFP;*Col-0 and *pSUC2:BRL3-GFP;brl3 roots* (Red box in f and g depicts the ROI used for measurements). Confocal images showing BRL3 expression pattern and vascular localization in **(k)** cotyledon and **(l)** root of *pSUC2:BRL3-GFP; brl3-2* transgenic seedlings. **(m)** Petiole elongation phenotypes and **(n)** quantification in WT (Col-0), *brl3-2* mutant and pSUC2:BRL3-GFP; brl3-2 transgenic plants grown at LD 22°C condition for 15-d and transferred to either 22°C or 28°C for 7-d. Boxplots depict the distribution of petiole lengths of rosette leaves from 22-d-old plants growing in control (dark green) or elevated temperature (light green) conditions. Different letters indicate a significant difference ($p\text{-value} < 0.05$) in a one-way ANOVA test plus Tukey's HSD test. Boxplot represent the median and interquartile range (IQR). Whiskers depict Q1 – 1.5*IQR and Q3 + 1.5*IQR and points experimental observations. Data from three independent biological replicates ($n > 30$).

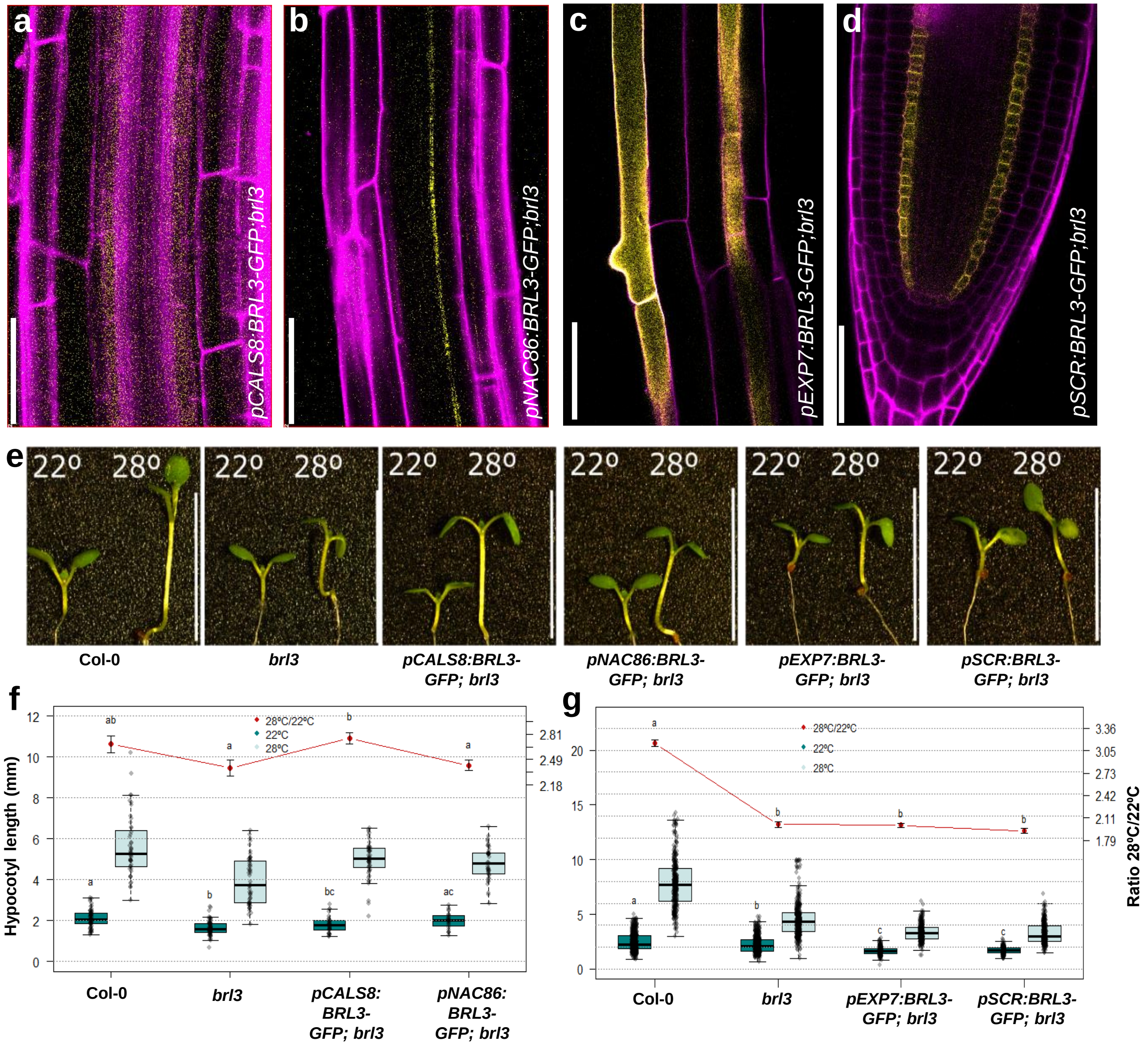


**Extended data figure S10. Local BRL3 expression in phloem can rescue *brl3* defects.**

Confocal imaging of 6-d-old **(a)** *pCALS8:BRL3-GFP;brl3* **(b)** *pNAC86:BRL3-GFP;brl3* **(c)** *pEXP7:BRL3-GFP;brl3* and **(d)** *pSCR:BRL3-GFP* seedling roots showing BRL3 expression in phloem pole pericycle, phloem sieve element, epidermis (trichoblasts) and endodermis respectively. **(e)** Pictures and (**f, g)** measurements of hypocotyl elongation response in WT (Col-0), *brl3* mutant, pCALS8:BRL3-GFP;brl3, pNAC86:BRL3-GFP;brl3, pEXP7:BRL3-GFP and pSCR:BRL3-GFP transgenic seedlings grown at 22 °C or 28 °C for 6 days under LD conditions. Scale bar: 10mm. Boxplots depict the distribution of hypocotyl lengths in control (dark green) or elevated temperature (light green) conditions. Boxplot represent the median and interquartile range (IQR). Whiskers depict Q1 – 1.5*IQR and Q3 + 1.5*IQR and points experimental observations. Red line depicts relative hypocotyl elongation upon high temperature (ratio 28 °C/22 °C ± s.e.m.). Data from three independent biological replicates (n > 50). Different letters represent significant differences (p-value < 0.05) in an ANOVA plus Tukey's HSD test.

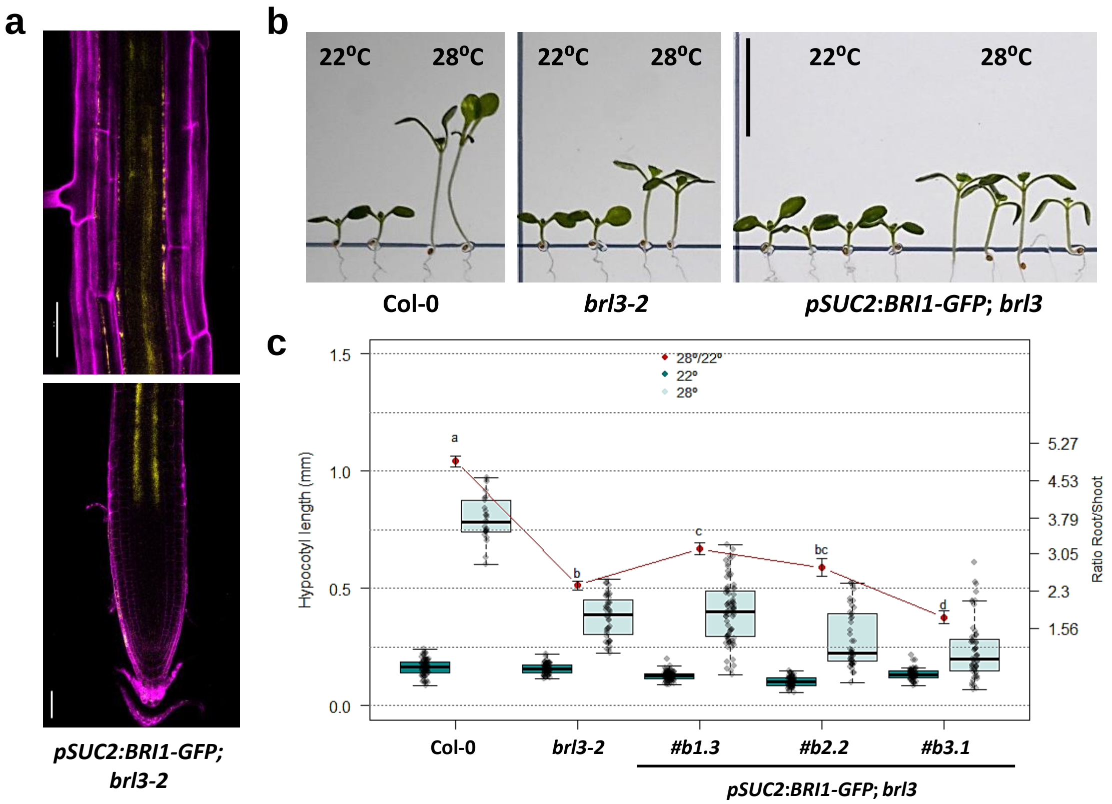


**Extended data figure S11. Local BRI1 expression from phloem companion cells can not rescue *brl3* thermomorphogenesis defects.**

**(a)**. Confocal imaging of primary roots maturation zone and meristematic regions of 7-d-old Arabidopsis *pSUC2:BRI1-GFP;brl3* seedlings. **(b)** Hypocotyl elongation phenotypes and **(c)** quantification in WT (Col-0), *brl3-2* mutant and *pSUC2:BRI1-GFP; brl3* transgenic grown at 22 °C or 28 °C for 6 days under LD conditions. Scale bar: 10mm. Boxplot represent the median and interquartile range (IQR). Whiskers depict Q1 – 1.5*IQR and Q3 + 1.5*IQR and points experimental observations. Red line depicts relative hypocotyl elongation upon high temperature (ratio 28 °C/22 °C ± s.e.m.). Data from three independent biological replicates (n > 30). Different letters represent significant differences (p-value < 0.05) in an ANOVA plus Tukey's HSD test.

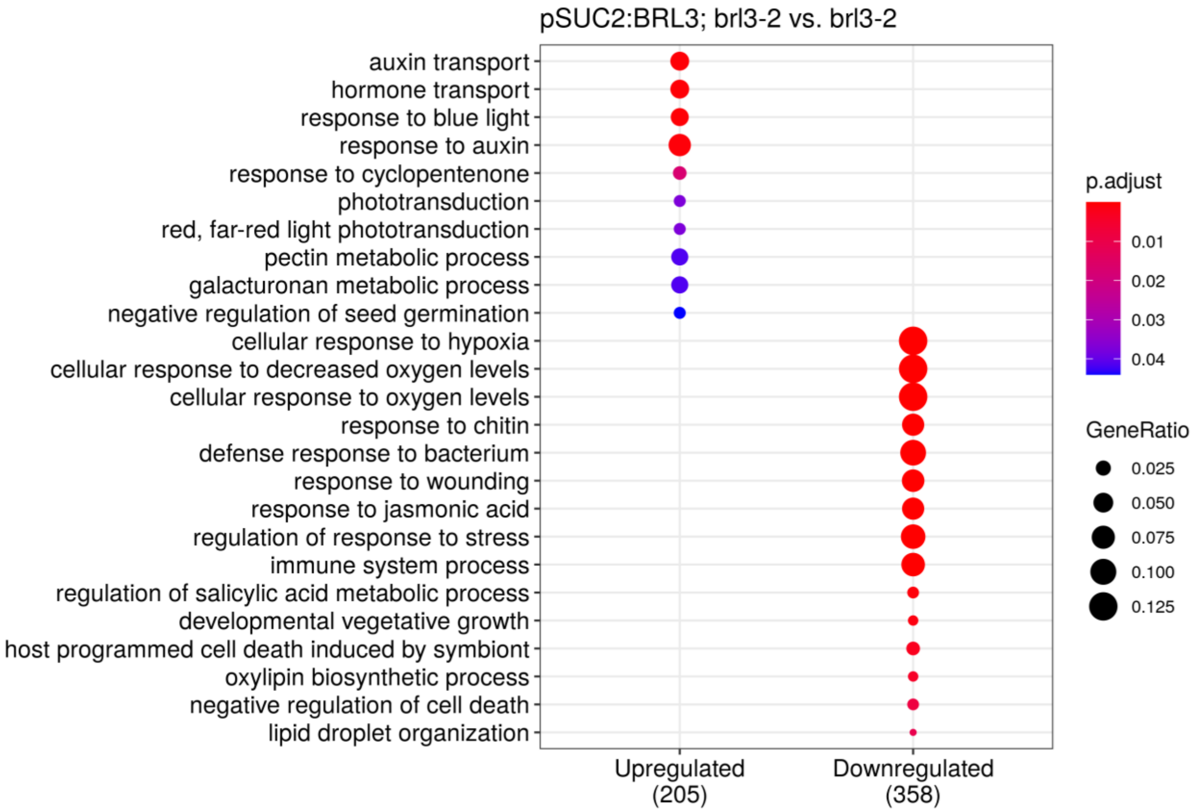


**Extended data figure S12. GO enrichment analysis of differentially regulated genes upon phloem specific BRL3 expression.**

GO enrichment analysis of genes up or downregulated in *pSUC2:BRL3-GFP; brl3-2* vs. *brl3-2* at ambient temperature (22°C) conditions. Size of the node represents the number of annotated genes in a particular category and the color indicates its adjusted p-value upon enrichment test.

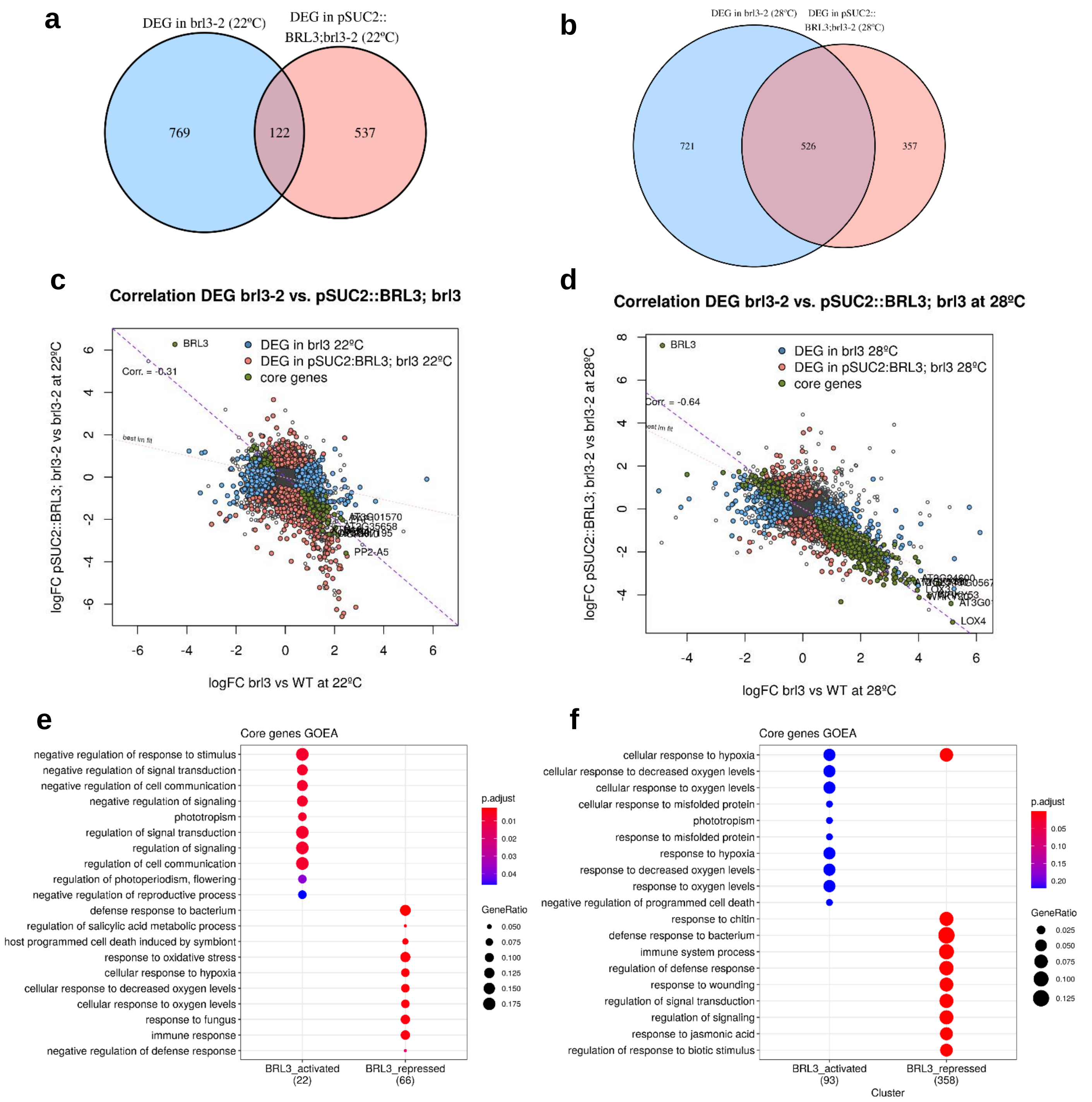


**Extended data figure S13. "Core Genes". Genes deregulated in both comparisons, *brl3* vs. WT and *pSUC2:BRL3; brl3* vs. *brl3,* and in opposite direction.**
**(a, b)** Overlapping of the transcriptional responses in the comparisons *brl3-2* vs. WT and pSUC2:BRL3; *brl3* vs. *brl3* at 22ºC (a) and 28ºC (b). Note the increased overlap at elevated temperature, up to 60% of DEG in *brl3*. **(c, d)** Scatter plots showing fold changes of both comparison (x-y axis) at 22ºC (c) and 28ºC (d). Genes following the inverse diagonal (purple) show an opposite behavior in *brl3* than in pSUC2:BRL3. These have high probabilities to be directly BRL3-regulated and we denominated them "Core genes". **(e, f)** GO enrichment analysis of the core genes, separating the enriched categories among the BRL3-activated genes (red) and BRL3-repressed genes (blue), either at 22ºC (e) and 28ºC (f).

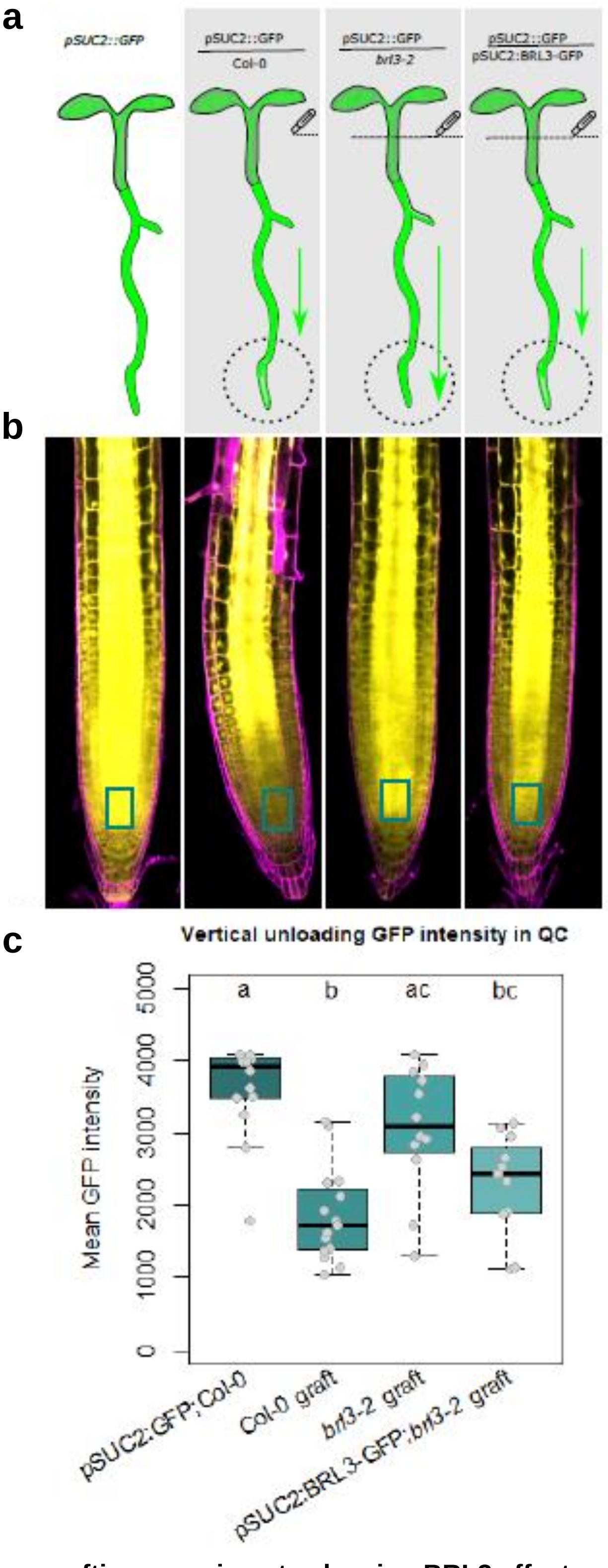


**Extended data figure S14. Micrografting experiments showing BRL3 effect on phloem function.**
**(a)** Graphical representation of the symplastic unloading of free-GFP using micrografting between *pSUC2::GFP* scion and rootstock from either WT Col-0 or *brl3* mutant seedlings. **(b)** Live confocal imaging of 6-day-post-grafting Arabidopsis root tips. Boxes account for the ROI area where GFP was measured, using the same ROI for all images placed right above the QC. In the labels, upper line indicates upper part of the chimeric plant after grafting (cotyledons and upper part of the hypocotyl), lower line indicates root part of the chimeric plant (upper part of the hypocotyl and primary root). **(c)** Mean GFP intensity in the selected area from B. Notice an increased unloading in root tip in brl3 root. Letters depict significant differences following ANOVA with Tukey Post-Hoc HSD test. data from three biological replicates, 9<n≤15.

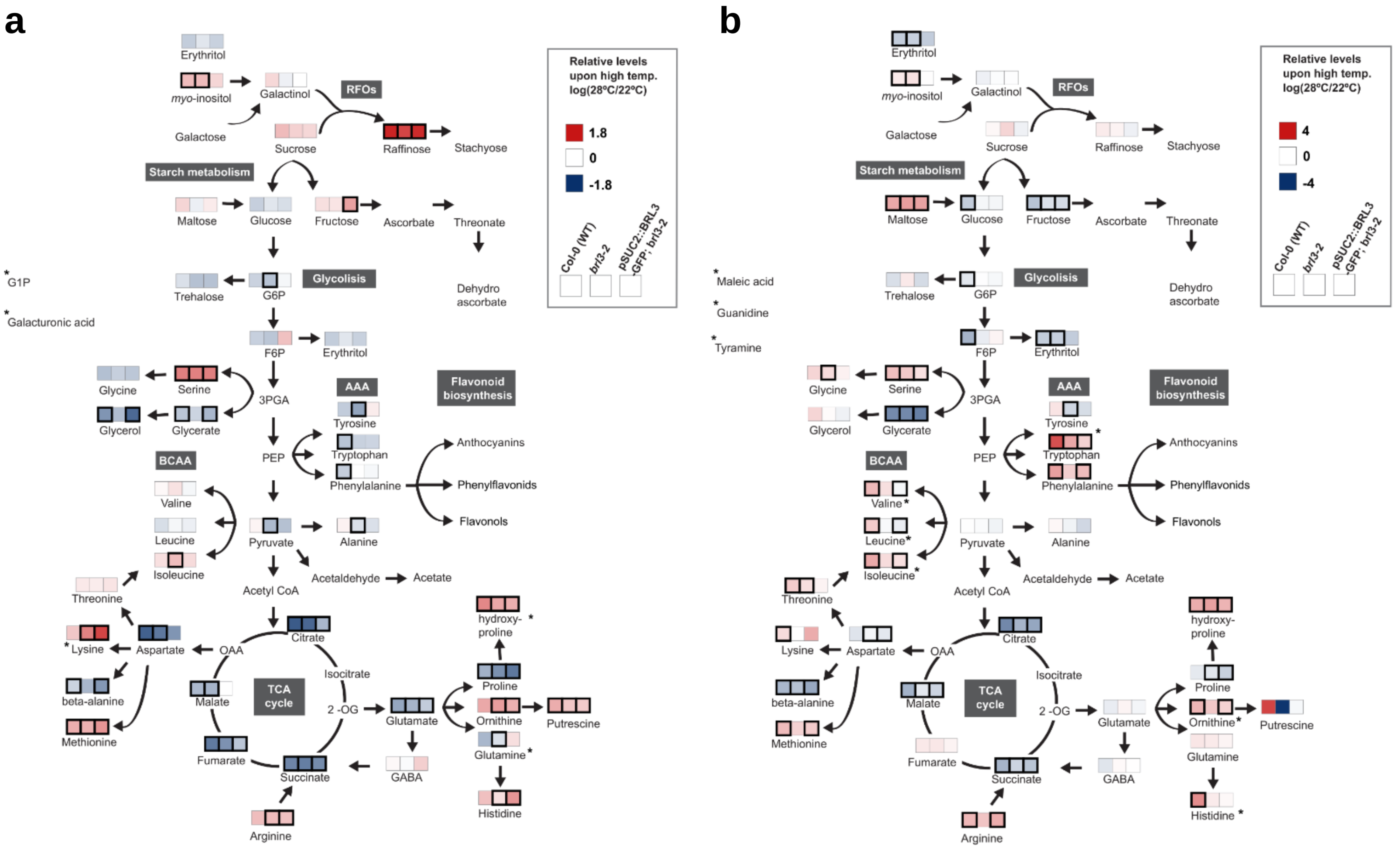


**Extended data figure S15. Metabolomics analysis of Col-0, *brl3* and pSUC2BRL3 under elevated temperatures**
Schematic representation of the specific metabolic response of **(a)** shoots and **(b)** roots of 6-d-old seedlings of WT (Col-0), brl3 and *pSUC2:BRL3-GFp;brl3* lines exposed to elevated temperature (28 °C) versus optimal temperature (28 °C) under LD conditions. Colored cells represent the log fold-changes. Statistically significant changes (28°C vs 22°C) are denoted by thick black borders on the boxes. Asterisks denote metabolites which have a statistically significant 28°C/22°C ratio between genotypes. Note that most of the metabolites respond similarly to temperature among genotypes.

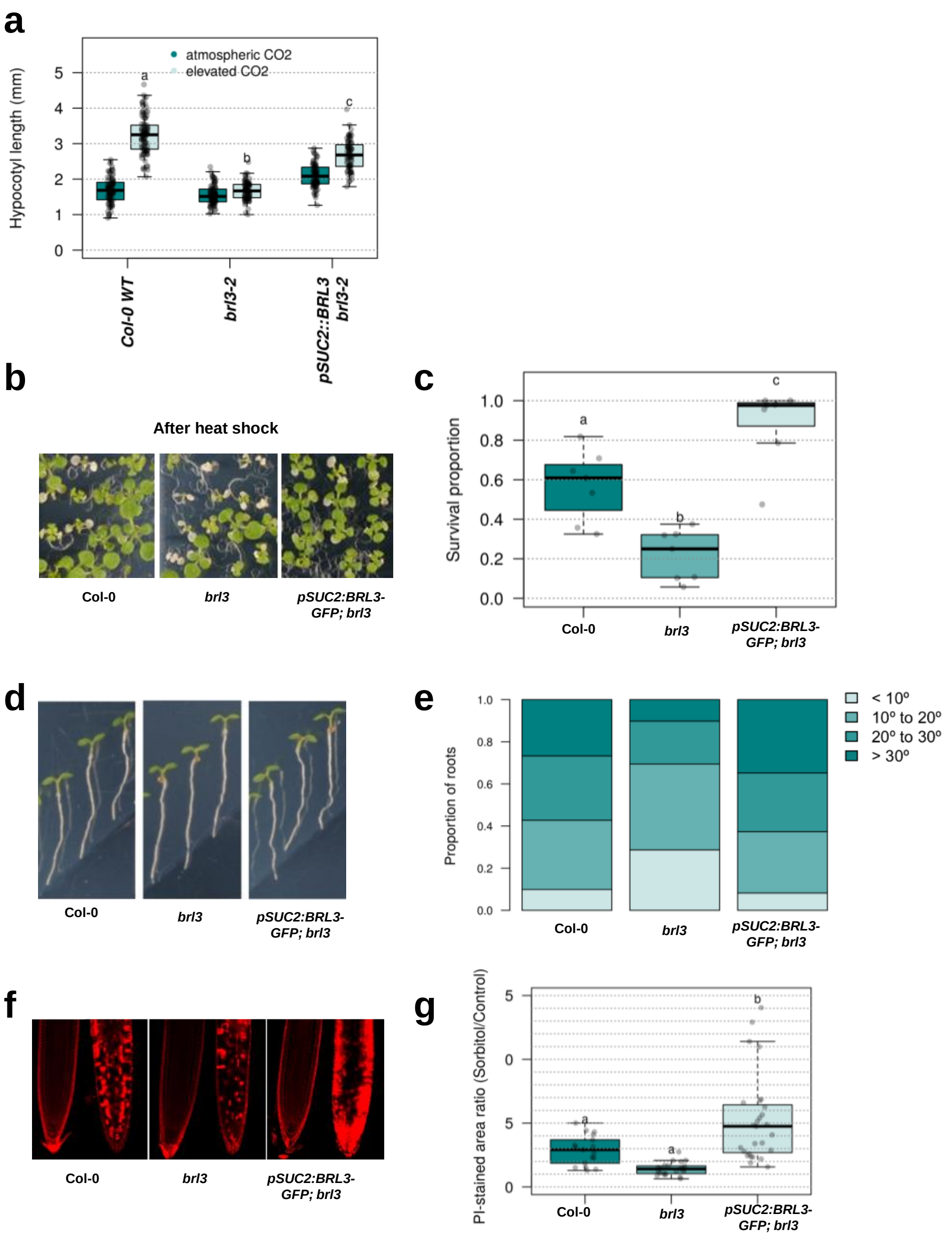


**Extended data figure S16. BRL3 signals from phloem regulate plant adaptation to multiple climate stress.**

**(a)** Graphs showing hypocotyl elongation growth response to elevated $CO_2$ in 6-d-old seedlings of WT, *brl3* and *pSUC2:BRL3-GFP;brl3* lines. **(b)** Pictures showing plant survival phenotypes after short-term heat-shock treatment (150 min at 42°C), **(c)** Quantification of survival rates upon heat-shock treatments. **(d)** Root hydrotropism response, **(e)** Distribution of the root angles upon hydrotropism response. **(f)** Cell death in root meristematic tissues after short term exposure to osmotic stress (8-10h), in WT, *brl3* and *pSUC2:BRL3-GFP;brl3* seedlings. **(g)** Quantification of cell damage due osmotic stress. As PI-stained ratio between treated and untreated roots. In all the conditions tested, phloem specific BRL3 expression was able to rescue stress responsive defects of the *brl3* mutant.
