## Supplementary material for "Brassinosteroid receptor BRL3 triggers systemic plant adaptation to elevated temperature from the phloem cells": Table S1. Primers used in this study

### EXTENDED DATA TABLES

#### **Table S1.** Primers used in this study

| Primer Id | sequence | purpose |
|---|---|---|
| bri4.2 | TTTAGGGTGAGCATGAGATCTCGTGGGCCG | genotyping *brl3-1* mutation |
| bri4.3 | GAAATCCCTGTAGGAATCGGAAAGCTTGAG | genotyping *brl3-1* mutation |
| JMRB | GCTCATGATCAGATTGTCGTTTCCCGCCTT | genotyping *brl3-1* mutation |
| SALK_005982_LP | ATATGGATGTTGCCGAATCTG | genotyping *brl1-2* mutation |
| SALK_005982_RP | CTGTAAAGCGCCATGACTAGC | genotyping *brl1-2* mutation |
| SALK_006024_LP | CCAGTGAACTCGTTTGAGCTC | genotyping *brl3-2* mutation |
| SALK_006024_RP | TTTATCGAACACTTTGTGGGC | genotyping *brl3-2* mutation |
| SALK_079612_LP | TCCCTAATTGCCAATCTTGAG | genotyping *brl3-3* mutation |
| SALK_079612_RP | CTGCACGAAAAGACCAAGAAG | genotyping *brl3-3* mutation |
| SALK_111696_LP | GCTTTACGCGAGTGTCTTGTC | genotyping *brl3-4* mutation |
| SALK_111696_RP | AGACAACAACCTTGTGGGATG | genotyping *brl3-4* mutation |
| LBb1.3 | ATTTTGCCGATTTCGGAAC | Universal T-DNA left border primer |
| bri1-301 F | GGAAACCATTGGGAAGATCA | genotyping *bri1-301* mutation[a] |
| bri1-301 R | GCTGTTTCACCCATCCAA | genotyping *bri1-301* mutation[a] |
| bes1-D F | TCGACGTCAGCTGCAGCT | genotyping *bes1-d* mutation[b] |
| bes1-D R | ATGGCTTAACTTGGCTGTTCT | genotyping *bes1-d* mutation[b] |
| M13 F | GTAAAACGACGGCCAG | Sequencing to check M13F region in a plasmid |
| AttR2/AttB2_Fw | GCTTTCTTGTACAAAGTGGT | Sequencing of the reporter in a plasmid |
| AttR2/AttB2_Rv | ACCACTTTGTACAAGAAAGC | Sequencing of the CDS insert in a plasmid |
| AtBRL3_gen_Rev | TCCGGAGGAGAAGTTGTTTC | Genotyping of transgenic plants |

[a] PCR product was digested by DpnII enzyme (R0543, New England Biolabs)

[b] PCR product was digested by MspI enzyme (R0106, New England Biolabs)
